# Non-linear effects of evening light exposure on cognitive performance

**DOI:** 10.1101/2025.03.16.643585

**Authors:** Amelie Reitmayer, Bilge Kobas, Michael Kammermeier, Carolina Rivera Luque, Kelly R Johnstone, Cassandra Madigan, Margaret M Cook, Thomas Auer, Siobhan Rockcastle, Manuel Spitschan

## Abstract

In humans, exposure to light can impact alertness and cognitive performance. These cognitive effects of light are mediated by the intrinsically photosensitive retinal ganglion cells (ipRGCs) expressing the photopigment melanopsin, which signals environmental light in addition to the cone- and rod-mediated pathways. Most studies investigating the cognitive effects of light have focused on alertness, raising the question how higher-level cognitive tasks such as working memory are modulated by light. This study investigated the dose-response relationship for alertness, cognitive performance and mental workload. Each level of melanopic illuminance (ranging from 1 lx to 595 lx melanopic EDI) was evaluated over separate days with a six-hour exposure in a controlled climate chamber with artificial lighting. Participants (n=16, 10 female, 27.4±2.5 years), completed the Psychomotor Vigilance Test (PVT) and *n*-back task every 30 minutes to assess reaction time, attention, and working memory, alongside subjective evaluations through questionnaires. The results suggest an inverted U-shaped correlation between cognitive functions and melanopic EDI and a U-shaped relationship between subjective assessments and melanopic EDI. Extreme lighting conditions in our stimulus set – both dim (1 lx melanopic EDI) and bright (595 lx melanopic EDI) – were associated with increased sleepiness and perceived workload, quicker reaction times, and diminished cognitive performance. Conversely, moderate illuminance levels (10 lx melanopic EDI and 70 lx melanopic EDI) positively influenced cognitive performance and mental workload but resulted in slower reaction times. This study illustrates that the relationship between melanopic EDI levels and cognitive performance does not follow a linear dose-response pattern, indicating a complex strategy for resource allocation in cognition.

## Introduction

Light exposure is key in regulating human physiology, influencing both circadian rhythms and cognitive function [1–3]. While extensive research has examined the alerting effects of light [2, 4–7], its impact on higher-level cognitive processes remains less well understood. The intrinsically photosensitive retinal ganglion cells (ipRGCs), which express the photopigment melanopsin, mediate non-visual responses to light [8, 9], including its effects on cognitive performance [4, 10]. Understanding how variations in melanopic equivalent daylight illuminance (melanopic EDI) influence cognitive function is critical for optimising indoor lighting environments.

Prior evidence has linked higher light levels to increased alertness [2, 4, 6]. Studies have shown that bright light exposure can enhance vigilance and reduce sleepiness, though its effects on more demanding cognitive tasks remain less clear [11–13]. The extent to which different lighting conditions optimise cognitive performance, particularly in indoor environments which can be crafted using targeted lighting design strategies [3, 14], warrants further investigation.

Research examining the impact of illuminance and temporal factors on cognitive performance and mental load has yielded mixed findings. Studies using the Psychomotor Vigilance Task (PVT) [15] suggest that under conditions of sleep deprivation, participants demonstrated faster reaction times and reduced sleepiness when exposed to bright illuminance (1000 lx) compared to dim lighting (<5 lx) [16]. However, another study found no significant differences in PVT reaction times between 1700 lx and 165 lx, although improvements were noted in cognitively demanding tasks such as the Backwards Digit-Span Task (BDST) at 1700 lx [11]. Notably, these studies did not identify any time-of-day effects on performance. Additional research supports the notion that higher illuminance levels enhance alertness compared to dimmer lighting conditions [17, 18]. Nighttime studies further indicate improved performance at 3000 lx relative to 100 lx, suggesting that this effect may be temporally constrained [19]. Similar findings regarding the influence of exposure duration have been observed during daytime hours [20].

This study examines the relationship between evening melanopic light exposure and cognitive performance, specifically focusing on reaction time, working memory, and subjective workload. Using a controlled experimental approach, participants were exposed to varying levels of melanopic EDI while completing cognitive tasks. By systematically assessing cognitive function across different lighting conditions, this study aims to refine our understanding of the effects of light on cognition and inform the design of lighting environments that better support cognitive performance.

### Objective

The objective of this study is the characterise the dose-response relationship between melanopic light exposure and alertness, cognitive performance and levels impact mental load (including momentary affect, perceived workload and sleepiness) during the evening.

## Methods

### Experimental design

#### Experimental setting

This study was conducted in August and September 2023 in the SenseLab, a dedicated custom-made testing space for simulating environmental conditions located at the Technical University of Munich (TUM). The SenseLab is equipped with a range of environmental sensors (see below under *Measurements, Environmental measurements*). The design is intended for various laboratory environment settings, accommodating configurations ranging from a single-user workspace through offices for up to three participants, to setups prioritising comfort and a calm atmosphere. It includes southeast-facing windows with adjustable blinds, ceiling fans, an air conditioning unit, and infrared heaters for effective thermal control.

#### Procedure

Participants were exposed to four different illuminance levels across four separate experimental sessions, each lasting five hours. The four lighting conditions are defined by logarithmic variations in melanopic illuminance, quantified using the melanopic equivalent daylight illuminance (melanopic EDI), given **Table 1**. Corneal-plane spectral irradiance distributions were measured using a spectroradiometer (Jeti spectraval 1511HiRes; Jena Technische Instrument GmbH, Jena, Germany). Irradiance spectra were then converted to photopic illuminance illuminance and melanopic EDI using the CIE-validated luox open-access platform [21]. Light was provided by overhead luminaires. The completed ENLIGHT Checklist [22] can be found in Supplementary Material S4.

**Table 1.** Vertical illuminance levels for each lighting condition and their associated melanopic EDI, as measured mean of the two seated positions.

| Lighting condition | Photopic illuminance at eye-level [lx] | Melanopic |
| --- | --- | --- |
| Very Dim | 1 lx vertical | 1 lx melanopic EDI |
| Dim | 15 lx vertical | 10 lx melanopic EDI |
| Moderately Bright | 90 lx vertical | 70 lx melanopic EDI |
| Very Bright | 800 lx vertical | 595 lx melanopic EDI |

An indoor operative temperature of 27°C were maintained throughout all measurement days. To ensure control over lighting conditions during daylight hours, the windows of the SenseLab were covered by opaque material to block out daylight. Every participant was scheduled to complete all four sessions within a maximum of five consecutive weeks. The end times of the sessions were adjusted according to their habitual bedtime that they indicated in their initial screening, which varied between 22:00 pm and 00:30 am. The sessions lasted five hours in the lab, and the participants were instructed to arrive six hours prior to the session start time.

At regular 30-minute intervals during each session, measurements were carried out to evaluate cognitive function and reaction time, along with subjective perceptions of mental load and sleepiness. This study is part of a larger experimental research project, including physiological measurements including salivary melatonin, core body temperature, and skin temperature, which will be analysed and presented in subsequent publications. In each session, two participants at the same time were situated in the SenseLab, one on the sofa and the other in the armchair (see **Figure S1**, **Supplementary Document S1**). Participants were asked to arrive one hour before the experiment’s onset to acclimate and consume a planned evening meal, adjusted according to their Body Mass Index (BMI). Throughout the session, consumption of food was not allowed. Participants were offered water every 30 minutes. A schematic outline of a single session’s structure is presented in **Figure 1**.

**Figure 1:**
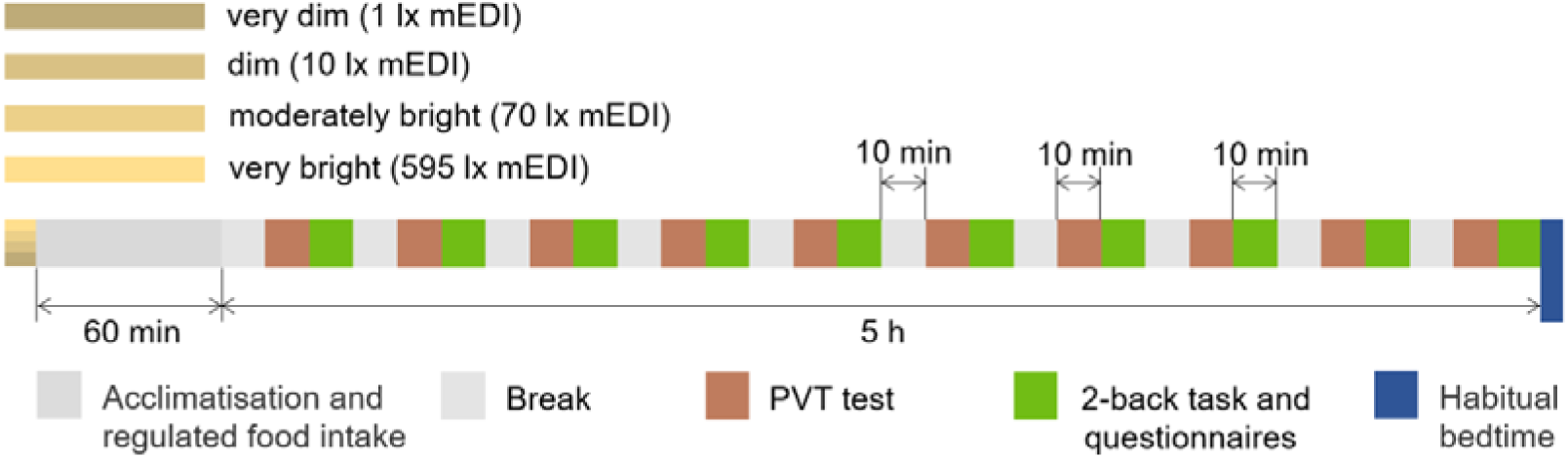
Experimental procedure replicated across all four sessions. In each session, a different illuminance level was introduced with the procedure remaining consistent. The sessions were started approx. 6 hours prior to the participants’ habitual bedtime.

#### Sample characteristics

A total of sixteen individuals participated in the study, with their demographic and physical characteristics presented in **Table 2**. Inclusion criteria required participants to be in good physical, mental, ocular, and retinal health, possessing normal visual acuity as determined by the Freiburg FrACTs sand Ishihara colour plates. Exclusion criteria were use of melanopic EDIcations, habitual smoking, engagement in shift work, and inter-time zone travel within the last three months. On each measurement evening, participants’ alcohol levels were verified to be 0‰ using a breathalyser (ACE AF-33; Ace Handels- und Entwicklungs GmbH, Freilassing, Germany). Participants were allowed to engage in activities such as using laptops or tablets, reading, or playing on handheld gaming devices, provided they remained seated. However, in the very dim (1 lx melanopic EDI) and dim (10 lx melanopic EDI) scenarios, looking at bright screens was restricted to avoid interference with the light conditions. If participants needed to leave the thermal chamber for bathroom breaks, they were required to wear special goggles that cut visible light transmission by 50% (Uvex Supravision Athletic). The required dress code for the participants included long pants, a short-sleeved shirt, sports shoes and normal short socks, ensuring a clothing factor of 0.5 based on ANSI/ASHRAE Standard 55 [23]. Participants were remunerated for each session they attended, with an additional bonus provided upon the completion of all four sessions. Detailed information about the study protocol and data handling procedures was shared with the participants and consent was obtained.

**Table 2.** Participant data (mean±1SD). BMI = Body mass index (Weight (kg)/[Height (m)]^2^).

|  | Males | Females | Total |
| --- | --- | --- | --- |
| **Sample size** | 6 | 10 | 16 |
| **Age (yrs.)** | 28.2±2.1 | 26.9±2.7 | 27.4±2.5 |
| **Height (cm)** | 178.5±8.0 | 167.3±6.8 | 171.5±8.9 |
| **Weight (kg)** | 78.4±15.2 | 68.9±15.1 | 72.4±15.4 |
| **BMI (kg/m<sup>2</sup>)** | 24.7±5.3 | 24.4±3.5 | 24.5±4.1 |

### Measurements

#### Environmental measurements

The climate chamber is equipped with a sensor kit, which contains several tinkerforge sensors to record environmental indoor parameters at a sampling rate of 60 seconds. The air temperature and globe temperature were recorded by Thermocouple Bricklets 2.0 with an accuracy of ±0.15%, relative humidity with the Humidity Bricklet 2.0 and an accuracy of ±2.0%, and CO_2_ concentration by the CO_2_ Bricklet 2.0 at an accuracy of ±30 ppm. The spectroradiometer Spectraval 1511 HiRes was used to measure the vertical illuminance. Participants were equipped with an ActTrust2 light sensor which was magnetically attached to the t-shirt at chest level to monitor light exposure.

#### Estimation of personalised light exposure

The ActTrust2 sensors were used to measure personal light exposure through an ambient light intensity sensor. The raw data from both sensors was combined, converted to photopic illuminance, and summarised every five minutes. To ensure accurate measurements, the first and last five minutes of each session were excluded to eliminate accidental light exposure when entering or leaving the lab. Additionally, the time spent outside the lab was removed by using the accelerometer data. To do this, the normalised activity value was calculated using the Proportional Integral Mode (PIMn) values, then transformed to a logarithmic scale. The top 10 percentile was considered as “walking” and used to indicate break-taking behaviour, which was subsequently filtered out. The remaining illuminance data was also log-transformed and controlled for any potential bias between the two experiment locations inside the lab (the sofa and armchair), as well as between the two sensors worn by different participants, using a mixed-effects model. The results of the model showed no significant effect for either location or individual sensor levels (see **Table 3**).

**Table 3.**
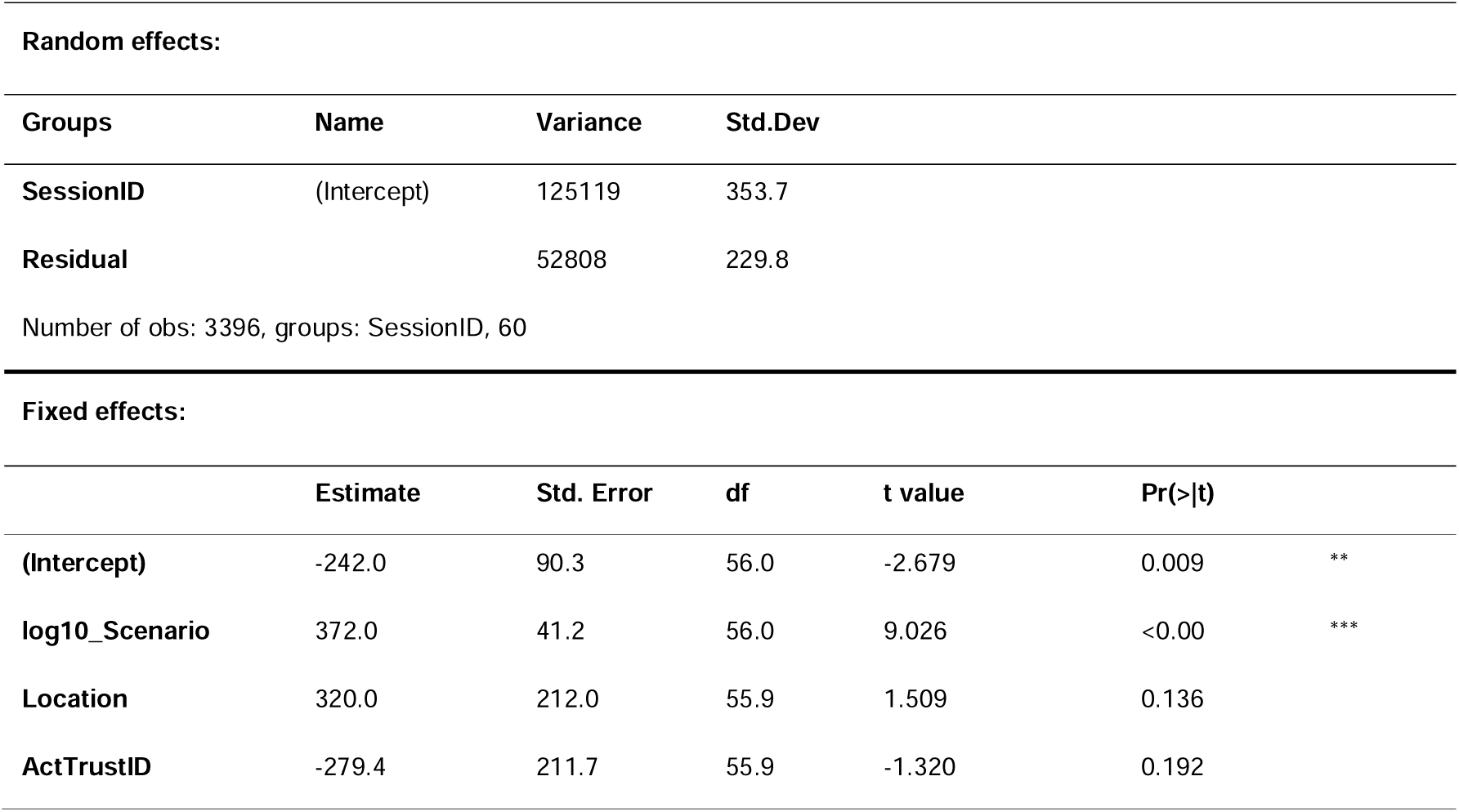
Mixed effects model for ambient light levels (log) against targeted light levels (log), different locations and different sensors.

| Random effects: |  |  |  |  |  |  |
| --- | --- | --- | --- | --- | --- | --- |
| Groups | Name | Variance | Std.Dev |  |  |  |
| SessionID | (Intercept) | 125119 | 353.7 |  |  |  |
| Residual |  | 52808 | 229.8 |  |  |  |
| Number of obs: 3396, groups: SessionID, 60 |  |  |  |  |  |  |
| Fixed effects: |  |  |  |  |  |  |
|  | Estimate | Std. Error | df | t value | Pr(>\|t\|) |  |
| (Intercept) | -242.0 | 90.3 | 56.0 | -2.679 | 0.009 | ** |
| log10_Scenario | 372.0 | 41.2 | 56.0 | 9.026 | <0.00 | *** |
| Location | 320.0 | 212.0 | 55.9 | 1.509 | 0.136 |  |
| ActTrustID | -279.4 | 211.7 | 55.9 | -1.320 | 0.192 |  |

#### Cognitive performance

##### Working memory-performance

Cognitive function, with focus on working memory, was measured using the n-back task [24]. The participants were presented with a sequence of letter stimuli. The n-back task was implemented via the web-based application PsyToolkit [25, 26] on a Samsung Galaxy Tab S7 Fe tablet with blue light filter, which was provided to the participants. For each stimulus, participants had to respond to whether the stimulus matched that one of n trials previously. The response had to be made within three seconds. There were 25 iterations of the trial for each test round, with the entire sequence being repeated twice.

The accuracy of responses, which was calculated using Equation (1), were used to evaluate the participants’ performance.

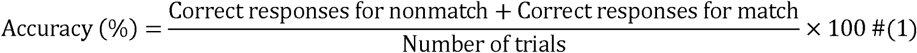

In this experiment, a 2-back condition was employed as the melanopic EDIum workload, a value which is considered the most reliable in terms of accuracy [27]. A correct response to a stimulus that matched its counterpart from two previous trials was recorded as a “match”, whereas correctness in cases where the participant withheld their response until a matching stimulus was presented was recorded as a “non-match”.

#### Alertness

Alertness was measured using the auditory psychomotor vigilance test (PVT) implemented on the PVT-192 Psychomotor Vigilance Task Monitor (Ambulatory Instruments, Inc., Ardsley, NY) [15]. During the 10-minute duration of the PVT, participants must react as quickly as possible to the appearance of visual stimuli in the form of red numbers on the small screen of the device. The stimuli appear randomly between two and ten seconds. Reaction time (RT) was used as the metric for evaluation. The PVT is highly reliable [28] and has a very low learning effect [29], and therefore is particularly suitable for retesting to examine the impact of light exposure on performance over time.

#### Subjective scales

All questionnaires on subjective perceptions were made accessible to each participant on the provided tablet.

The NASA-TLX (National Aeronautics and Space Administration Task Load Index), a prevalent method for assessing workload [30, 31], was employed for participants’ self-evaluation of their workload. In this assessment, participants evaluated their mental and temporal workload, along with subjective performance, experienced during the cognitive function and reaction time tests. These evaluations were quantitatively recorded on a 20-point scale. Participants were instructed to complete the NASA-TLX questionnaire immediately after the cognitive tasks to accurately capture their perceptions during these tasks.

The Momentary Affect Scale (MAS), known for its sensitivity in measuring affective states at consecutive time intervals [32], was used. This instrument allowed participants to self-report their levels of energetic and tense arousal at the measurement time points on an 11-point scale.

The validated and reliable Karolinska Sleepiness Scale (KSS) [33] was employed to assess subjective sleepiness. Through this scale, participants rated their sleepiness on a 9-point scale, providing insights into their subjective state of alertness.

### Statistical analyses

The statistical analysis of the single variables was performed using the R statistical computing environment (version 4.3.0 [34]). Results of the lighting scenarios, which were treated as multiple paired groups, were tested for normality using Shapiro-Wilk test. Following this, an ANOVA test was performed to examine differences among the four groups. If significant differences were found, Tukey’s HSD test was used for conducting multiple comparisons between the groups. The significance of differences is considered at p ≤ 0.05. Outliers were removed using the Interquartile Range (IQR) method with a coefficient of 1.5. A linear mixed model (LMM) was employed, with scenarios and time points designated as fixed effects, to examine their impact on the outcome measures, which were modelled as dependent variables. Participants were treated as random effects to account for interindividual variability. The LMM approach was chosen for its ability to accurately estimate complex interactions between variables. To stabilise the exponential variance in the illuminance values, a log transformation was applied to the scenarios prior to the analysis. In the analysis of the relationship between illuminance scenarios and the outcome variables, both linear and quadratic terms of the scenarios were incorporated to enable the capture of non-linear relationships. To reduce the risk of type I errors associated with multiple comparisons, Bonferroni correction was included to improve the statistical accuracy.

#### Data exclusion

Prior to the data analysis, the data was screened for technical errors and protocol deviations. Participant #103 was excluded from all analyses due to a discrepancy in the personal ambient illuminance measurements following an error in experimental condition assignment.

#### Ethics approval

The study received ethical approval from the TUM Ethics Committee (2023-311-S-KH).

## Results

The analysis of the measurements is organised into three parts: results from objective measures like cognitive performance, results from subjective measures such as perceptions of mental load and sleepiness, and the environmental conditions. All results from the Linear Mixed Model (LMM) analysis are included in the supplementary material.

### Alertness and cognitive performance

Analysis of the psychomotor vigilance test (PVT) results showed that exposure to different melanopic EDI levels affects reaction and attention. **Figure 2** shows the variation in reaction times across the four illuminance scenarios and the timepoint of the measurement throughout the experimental session. The shortest reaction time was observed in the very bright (595 lx melanopic EDI) condition. However, the data did not exhibit a linear dose-response relationship as illuminance levels decreased. Reaction times were adversely affected at dim (10 lx melanopic EDI) and moderately bright (70 lx melanopic EDI) levels but showed improvement at the very dim (1 lx melanopic EDI) level. Pairwise comparisons revealed significant differences (p < 0.01) between the very bright (595 lx melanopic EDI) scenario and both dim (10 lx melanopic EDI) and moderately bright (70 lx melanopic EDI) scenarios as well as between the very dim (1 lx melanopic EDI) scenario and both dim (10 lx melanopic EDI) and moderately bright (70 lx melanopic EDI) scenarios.

**Figure 2.**
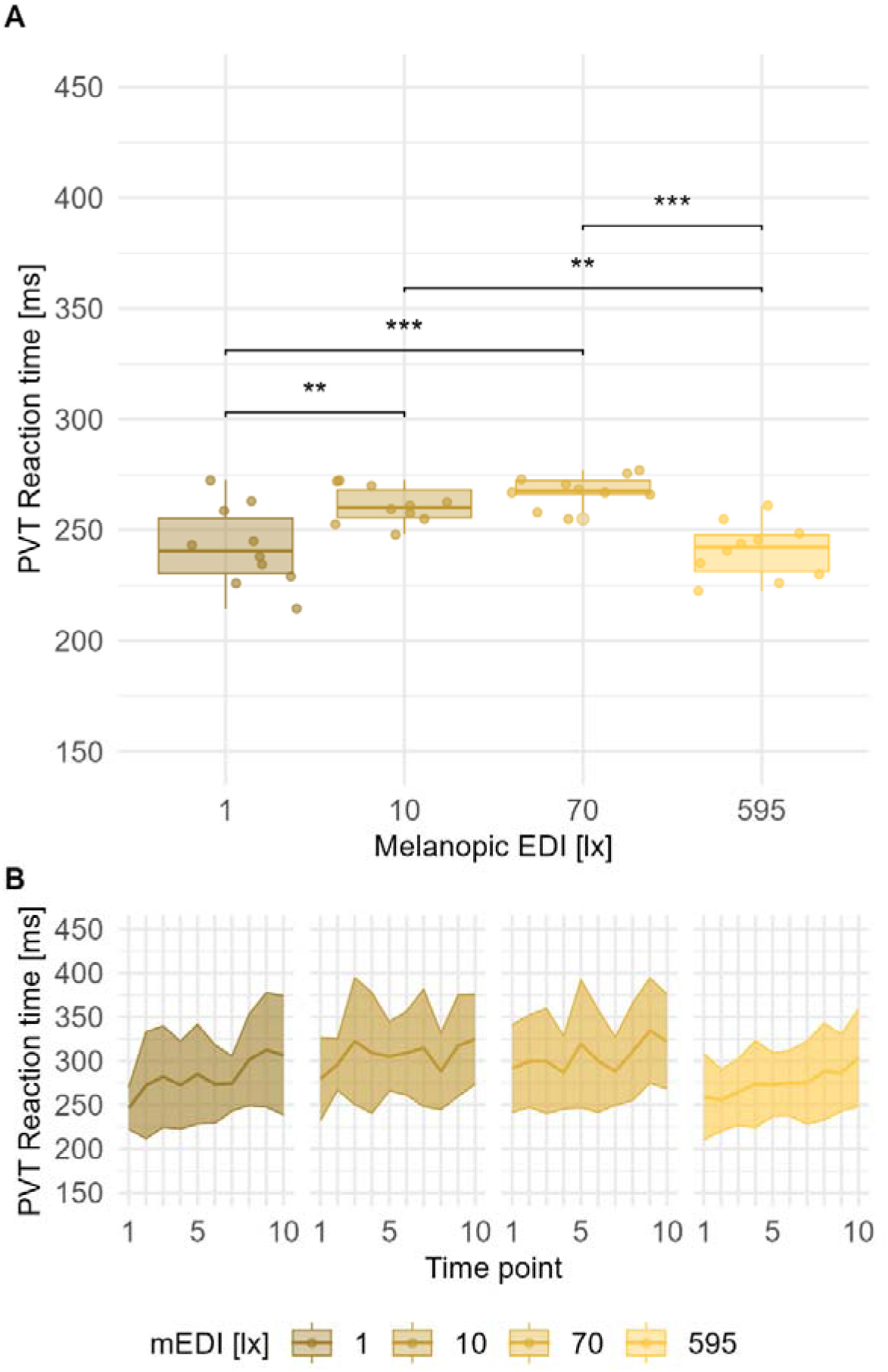
PVT Reaction time by light scenario (A) and time of the measurements (B) with the symbols in (A) indicating the following statistical significances: ns: p > 0.05; *: p ≤ 0.05; **: p ≤ 0.01; ***: p ≤ 0.001; ****: p ≤ 0.0001

Furthermore, Linear Mixed Model (LMM) analysis also indicated the presence of an inverted U-shaped relationship, as indicated in the graph. The significant (p < 0.001) results of the LMM analysis suggest that first, reaction times lengthen with increasing melanopic EDI until a certain level. Beyond that certain melanopic EDI level, the reaction time lowers again until reaching the brightest melanopic EDI level. A significant relationship (p < 0.001) was also observed between the reaction time and the measurement time, indicating that the reaction time deteriorates over the course of the evening across all scenarios.

Similarly to the reaction and attention of the participants, working memory was also affected by different melanopic EDI levels. **Figure 3** shows working memory performance, as indicated by the accuracy level of the performed n-back task. Here, a non-linear relationship formed as an inverted U-shape can be observed. The highest accuracy was observed at the moderately bright (70 lx melanopic EDI) scenario, showing significant differences when compared to very bright (595 lx melanopic EDI) (p < 0.05), and to very dim (1 lx melanopic EDI) (p < 0.00), with very dim (1 lx melanopic EDI) demonstrating significantly lower accuracy than all other scenarios. This observation is underlined by the linear mixed model (LMM) analysis (**Table S1**, **Supplementary Document S3**) showing that, up to a certain point, the accuracy increases with melanopic EDI. However, this trend starts to reverse above a certain melanopic EDI level, which can be identified as moderately bright (70 lx melanopic EDI) illuminance (**Figure 3**).

**Figure 3:**
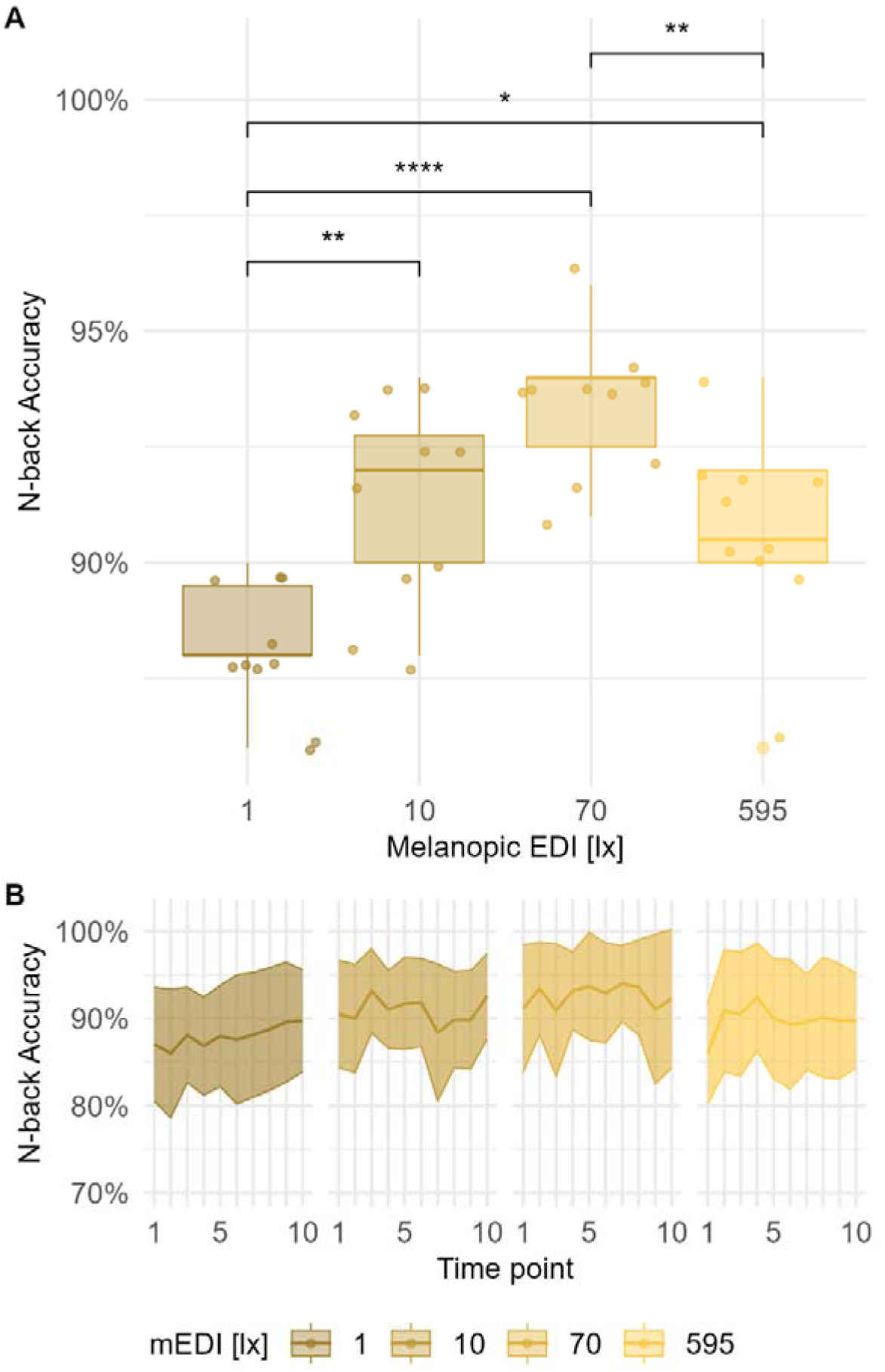
N-back Accuracy by light scenario (A) and time of the measurements (B) with the symbols in (A) indicating the following statistical significances: ns: p > 0.05; *: p ≤ 0.05; **: p ≤ 0.01; ***: p ≤ 0.001; ****: p ≤ 0.0001.

### Subjective scales

Results for perceived workload and momentary affect states show significant interactions with changing melanopic EDI. **Figure 4**, **Figure 5** and **Figure 6** provide insights into subjective feelings experienced during cognitive performance tasks, focusing on mental and temporal demand as well as self-rated performance by the participants. Similar to the accuracy findings in the performance task, these figures also illustrate a U-shaped pattern, with the moderately bright (70 lx melanopic EDI) scenario showing the lowest perceived load and highest performance rating. Both mental and temporal demand show a significant increase (p < 0.01) moving away from the moderately bright (70 lx melanopic EDI) level, with the highest loads reported in the very dim (1 lx melanopic EDI) scenario. Differences in self-estimated performance among the very bright (595 lx melanopic EDI), dim (10 lx melanopic EDI), and very dim (1 lx melanopic EDI) scenarios do not reach statistical significance (see **Figure 6**). Results from the LMM analysis (detailed in **Table S1**, **Supplementary Document S3**) show similar trends and significances across the scenarios (p < 0.00), indicating the lowest mental and temporal demand at moderate melanopic EDI levels and higher demands in both lower and brighter melanopic EDI levels. **Table S1** (**Supplementary Document S3**) also presents a significant temporal dependency (p < 0.05) in temporal demand that indicates a decline in demand over time, especially pronounced in the lowest melanopic EDI exposure. However, the self-rated performance of the participants showed neither a non-linear relationship with illuminance levels nor a significant correlation with the timepoints of the measurements.

**Figure 4:**
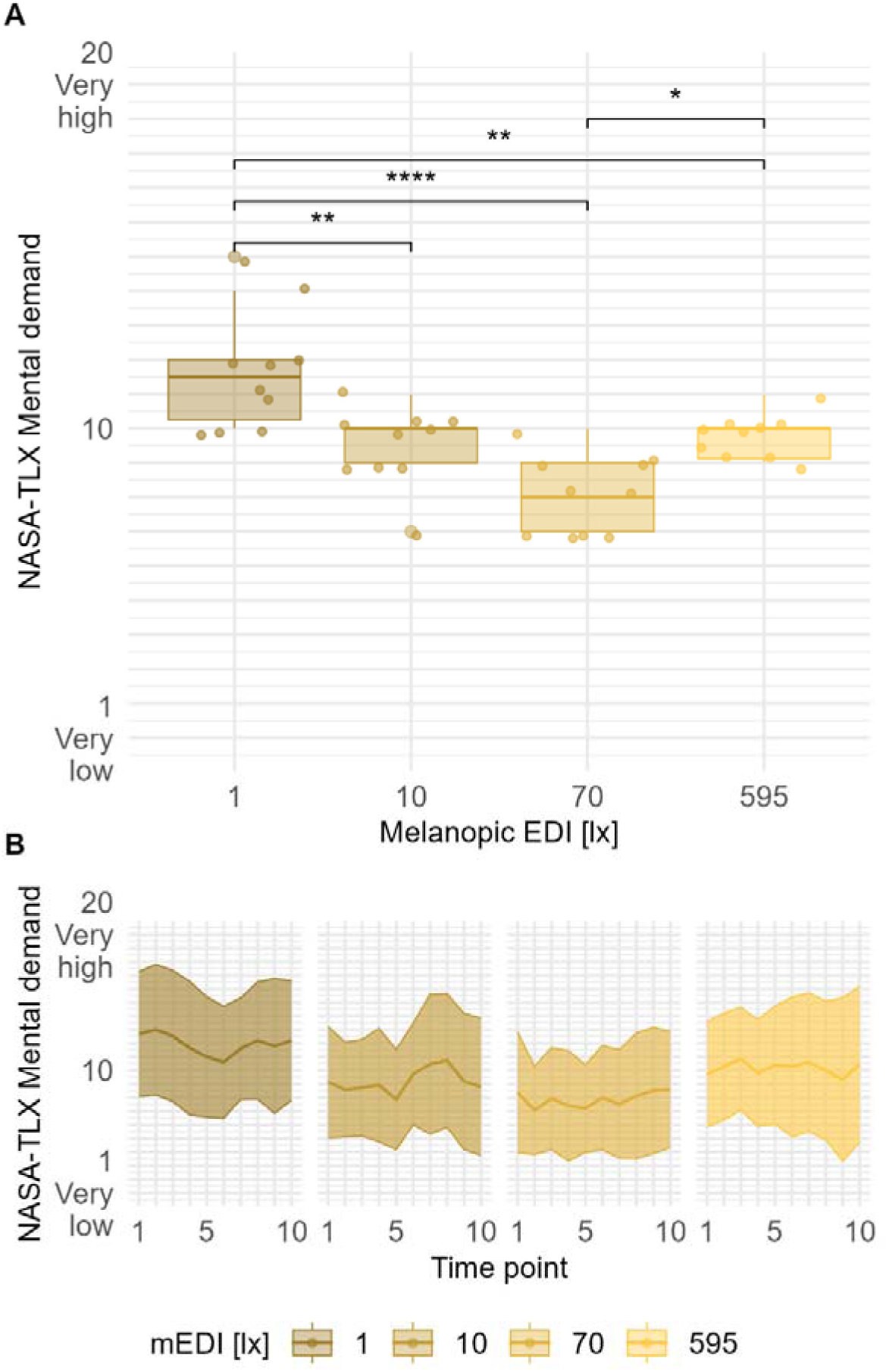
NASA-TLX Mental demand by light scenario (A) and time of the measurements (B) with the symbols in (A) indicating the following statistical significances: ns: p > 0.05; *: p ≤ 0.05; **: p ≤ 0.01; ***: p ≤ 0.001; ****: p ≤ 0.0001

**Figure 5:**
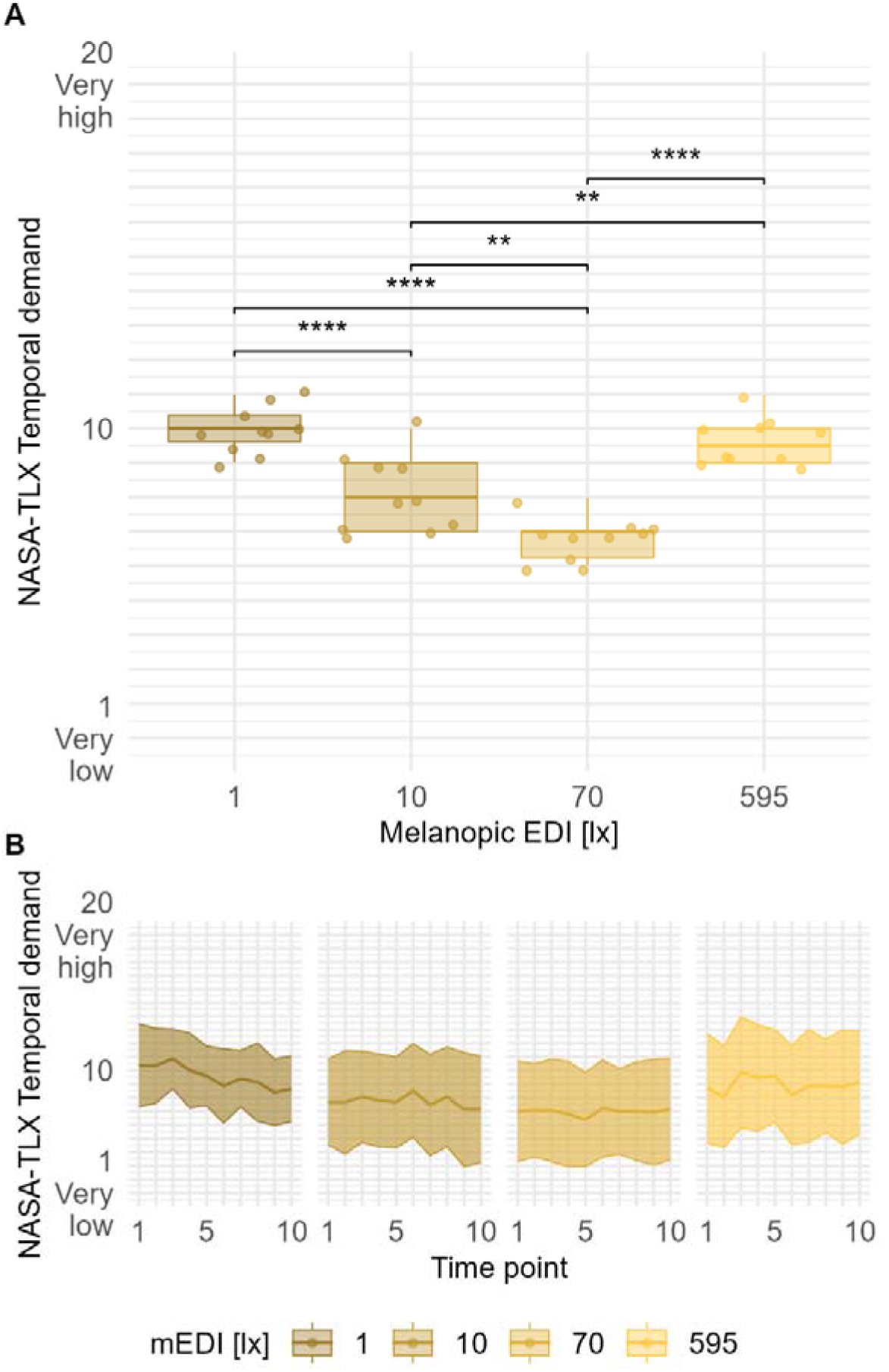
NASA-TLX Temporal demand by light scenario (A) and time of the measurements (B) with the symbols in (A) indicating the following statistical significances: ns: p > 0.05; *: p ≤ 0.05; **: p ≤ 0.01; ***: p ≤ 0.001; ****: p ≤ 0.0001

**Figure 6:**
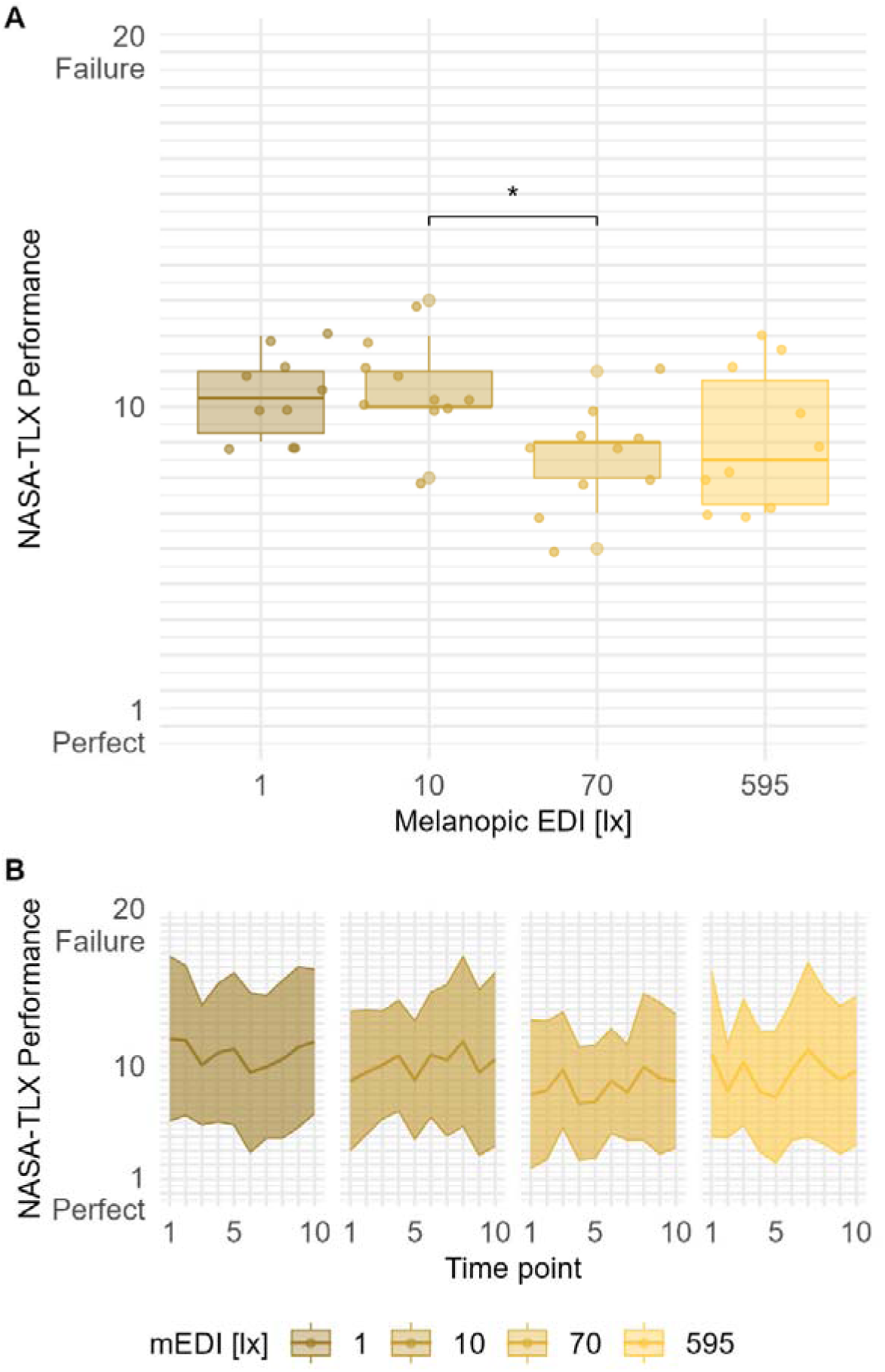
NASA-TLX Performance by light scenario (A) and time of the measurements (B) with the symbols in (A) indicating the following statistical significances: ns: p > 0.05; *: p ≤ 0.05; **: p ≤ 0.01; ***: p ≤ 0.001; ****: p ≤ 0.0001

Similarly to the mental workload responses, participants expressed different momentary affect states depending on the melanopic EDI levels. The pairwise comparisons, as depicted in **Figure 7** and **Figure 8**, show minimal significant differences in arousal levels among the scenarios. However, LMM analysis indicates a U-shaped relationship between the scenarios and tense arousal (**Table S1, Supplementary Document S3**). While perceived stress levels and nervousness tend to decrease with increasing light intensity, this tendency decreases from a certain melanopic light intensity and reaches reverse statements in the brightest melanopic EDI exposure. Specifically, participants reported feeling most relaxed in the moderately bright (70 lx melanopic EDI) scenario and most dull in the dim (1 lx melanopic EDI) scenario. Energetic arousal was observed to increase linearly with illuminance, reaching a maximum at the very bright (595 lx melanopic EDI) level. Furthermore, the analysis of energetic arousal reveals a significant temporal trend, showing a consistent decline in arousal levels across all scenarios as the evening progressed.

**Figure 7:**
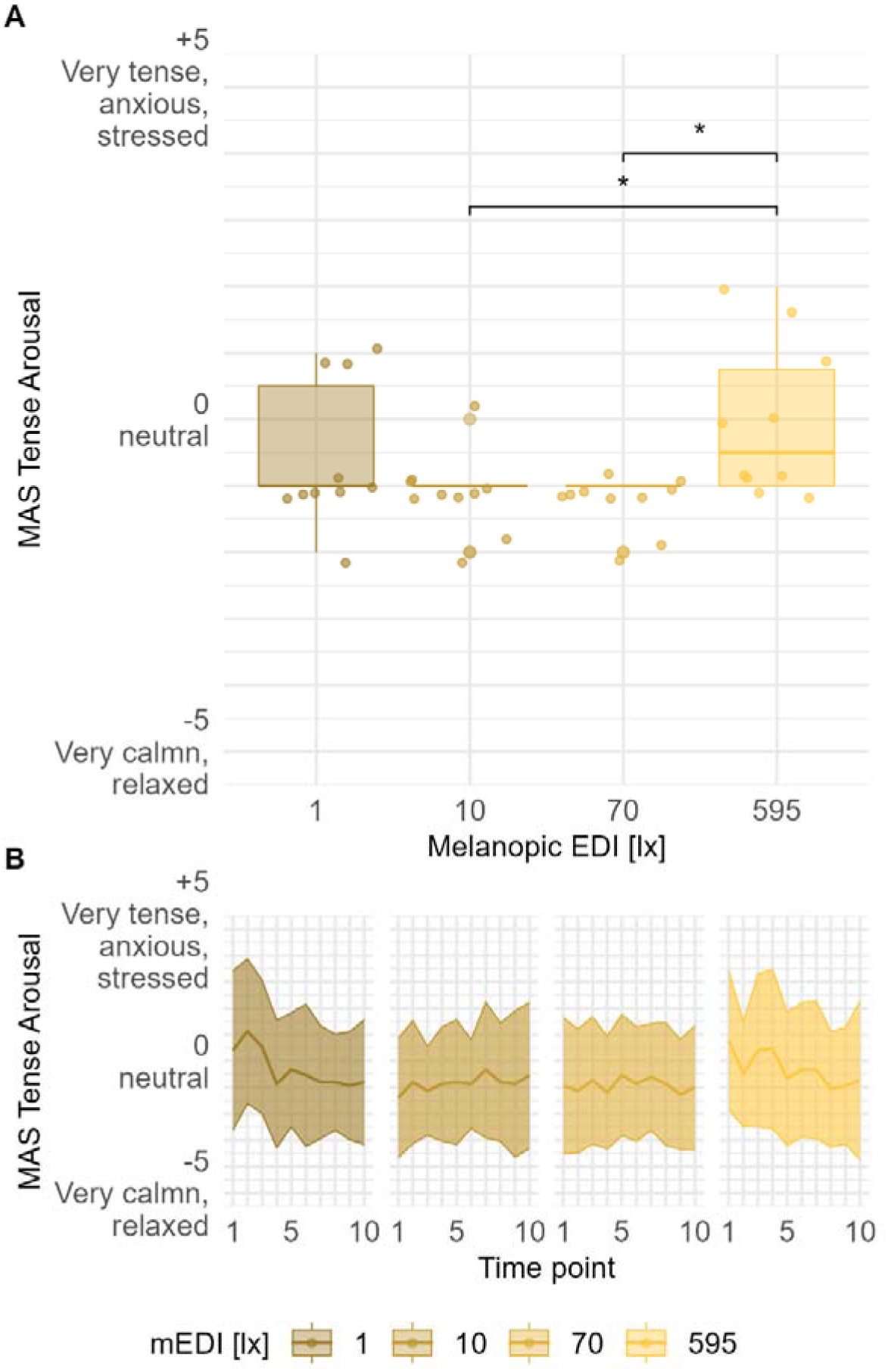
MAS Calmness/ Tense arousal by light scenario (A) and time of the measurements (B) with the symbols in (A) indicating the following statistical significances: ns: p > 0.05; *: p ≤ 0.05; **: p ≤ 0.01; ***: p ≤ 0.001; ****: p ≤ 0.0001

**Figure 8:**
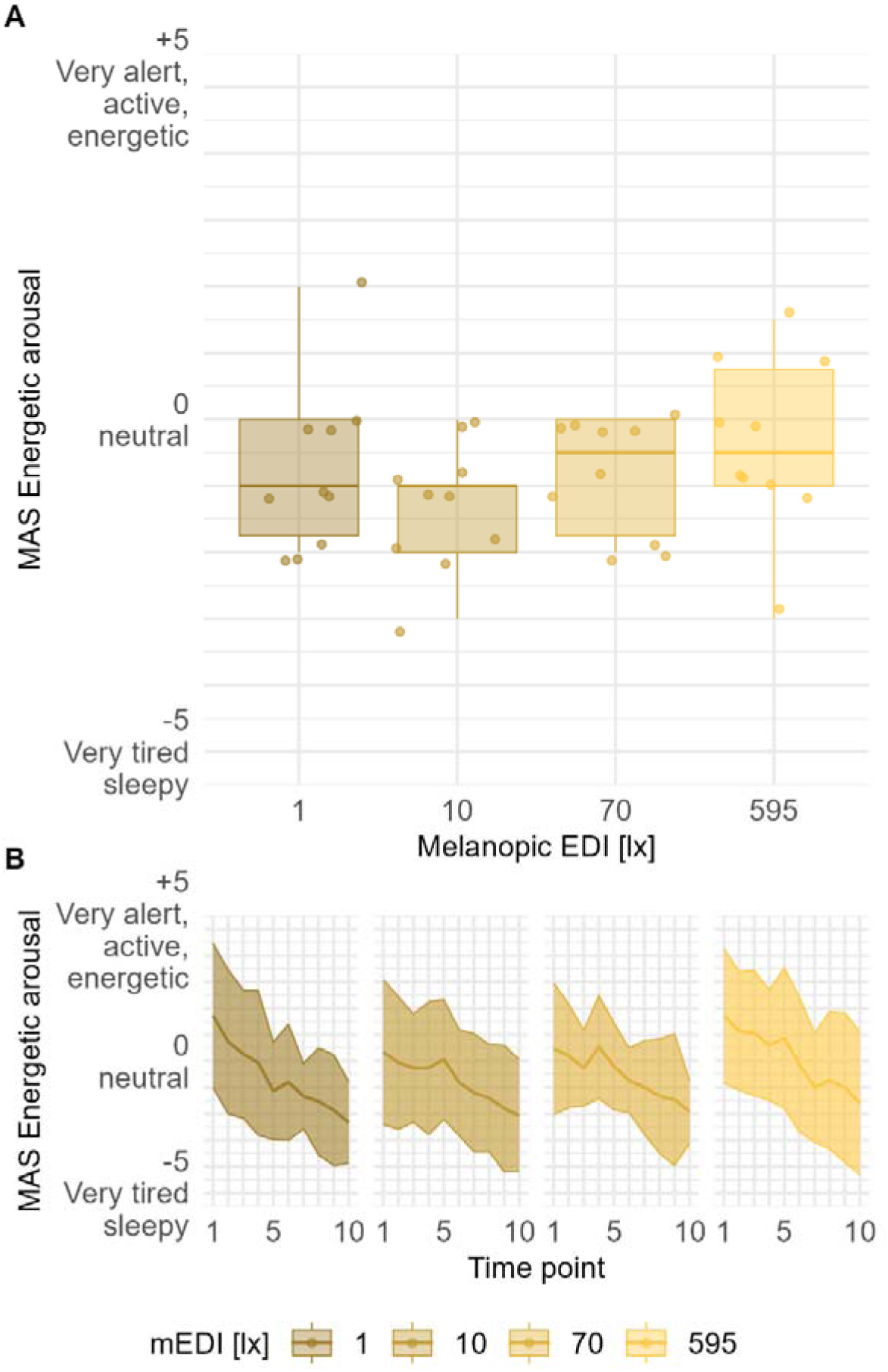
MAS Energetic arousal by light scenario (A) and time of the measurements (B) with the symbols in (A) indicating the following statistical significances: ns: p > 0.05; *: p ≤ 0.05; **: p ≤ 0.01; ***: p ≤ 0.001; ****: p ≤ 0.0001

The alertness of the participants was significantly affected by the different melanopic EDI levels. **Figure 9** illustrates subjective sleepiness levels, showing that participants’ state of alertness was significantly lower in the very dim (1 lx melanopic EDI) scenario than in scenarios with higher illuminance levels (p < 0.05). Furthermore, a positive linear relationship with the time of measurement was observed across all scenarios. The LMM analysis (**Table S1**, **Supplementary Document S3**) showed that sleepiness intensified as measurements were taken later in the session (p < 0.00). Separately, the analysis indicates a U-shaped trend with increasing melanopic EDI levels, alertness was rated higher (p < 0.00) and at the highest, very bright (595 lx melanopic EDI) level, sleepiness increased. Notably, in the moderately bright (70 lx melanopic EDI) scenario, participants felt the least sleepy.

**Figure 9:**
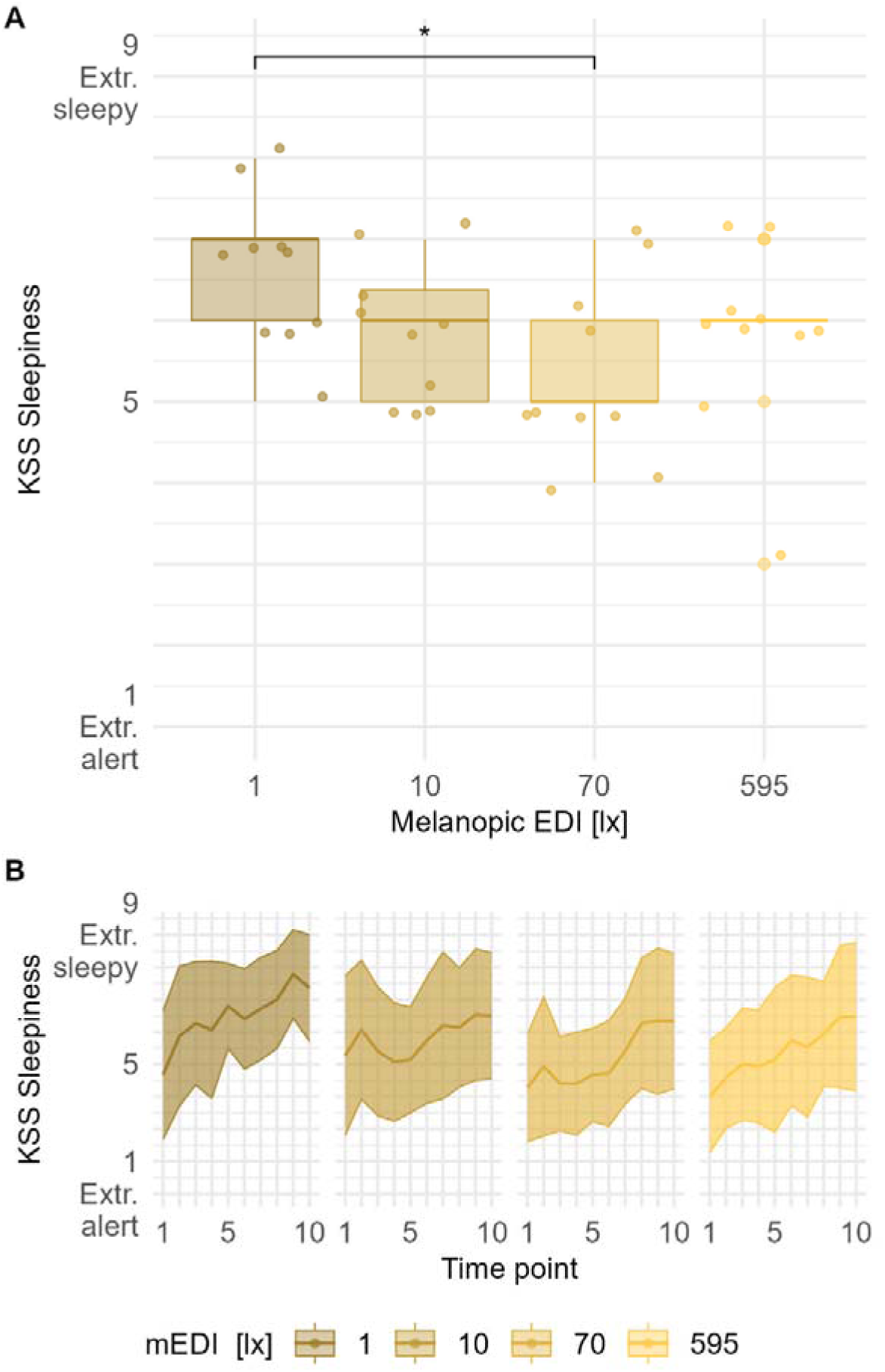
Karolinska sleepiness scale votes by light scenario (A) and time of the measurements (B) with the symbols in (A) indicating the following statistical significances: ns: p > 0.05; *: p ≤ 0.05; **: p ≤ 0.01; ***: p ≤ 0.001; ****: p ≤ 0.0001

#### Environmental measurements

**Table 4** summarises the environmental conditions for the four illuminance scenarios investigated. The melanopic EDI levels were kept constant throughout the day and each session. The sensor box was placed at the height of the sitting level in between the two participants at ca. 100 cm from each participant.

**Table 4:** Indoor environmental parameters for the combined data of all sessions (mean±1SD)

| Scenario<br>(lx<br>melanopic<br>EDI) | Measured<br>illuminance (lx) | Air<br>temperature<br>(°C) | Globe<br>temperature<br>(°C) | Operative<br>temperature<br>(°C) | Relative<br>humidity (%) | CO<sub>2</sub> level (ppm) |
| --- | --- | --- | --- | --- | --- | --- |
| 1 | 1.40 ± 10.77 | 27.60 ± 0.60 | 26.48 ± 0.54 | 27.05 ± 0.57 | 52.01 ± 5.51 | 564.95 ± 103.84 |
| 10 | 14.96 ± 19.25 | 27.48 ± 0.64 | 26.41 ± 0.53 | 26.95 ± 0.59 | 51.19 ± 4.84 | 591.81 ± 76.05 |
| 70 | 123.61 ± 65.83 | 27.39 ± 1.10 | 26.41 ± 0.50 | 26.90 ± 0.57 | 51.66 ± 6.93 | 574.94 ± 101.79 |
| 595 | 1168.44 ± 650.85 | 27.74 ± 0.75 | 26.89 ± 0.58 | 27.32 ± 0.67 | 50.43 ± 5.50 | 595.99 ± 149.26 |

## Discussion

This study investigated how different levels of illuminance (very dim, dim, moderately bright and very bright) and their corresponding non-visual effects in melanopic Equivalent Daylight Illuminance (melanopic EDI)=:J (1 lx, 10 lx, 70 lx, 595 lx) affect cognitive performance and mental load, which comprises momentary affect states and perceived workload. Additionally, it assessed how these relationships are impacted by temporal effects, including the time of day and exposure duration. The research was conducted in four separate sessions, each assessing one illuminance level, beginning at 4.00 pm earliest, and spanning six hours, with the arrival and departure time coordinated with the participants’ habitual bedtime. Every 30 minutes, participants engaged in Psychomotor Vigilance Task (PVT) and 2-back tasks, during which assessments of mental load were also collected. To better capture interaction effects of the illuminance and temperature, these environmental parameters were maintained in a constant intensity and level throughout each session.

The findings revealed an inverted U-shaped relationship between melanopic EDI levels and cognitive performance. This pattern was corroborated by participants’ self-reports of temporal and mental demand as well as alertness, which showed a matching U-shaped relationship. However, this pertains specifically to performance on the 2-back task, where higher accuracy was associated with lower perceived workload. In contrast, the PVT outcomes presented a paradox, showing the quickest reaction times under conditions of highest perceived workload. These results diverge from previous studies that identified a positive relationship between illuminance and performance, [16, 35], where typically only two illuminance levels were examined, limiting the ability to find a non-linear or specific dose-response relationship. The differences in reactions to the PVT and n-back tasks align with findings from other research [36, 37], pointing out that cognitive functions during n-back tasks improved under 200 lx compared to 1000 lx, while brighter illuminance of 1000 lx enhances reaction times in PVT tests. The findings suggest that participants show quicker reactions under greater mental and temporal stress, whereas higher cognitive functioning is observed under reduced perceived workload. Melanopic EDI levels which are either very dim (1 lx melanopic EDI) or very bright (595 lx melanopic EDI) appeared to trigger an increase in mental and temporal load, whereas moderate melanopic EDI levels (10 lx melanopic EDI and 70 lx), were associated with a lower perceived workload. At the same time participants were most alert at melanopic EDI of 70 lx, measured by their energetic arousal state and felt sleepiest at melanopic EDI of 1 lx and 10 lx. The lack of a clear correlation due to the inverted U-shape in PVT complicates the understanding of the dynamics between alertness and reaction time. The observed inverted U-shape between cognitive functions and log-transformed melanopic EDI challenges previously established relationships, suggesting that the impact of photopic illuminance on alertness and cognitive performance may vary throughout the day and under different conditions. A dose-response relationship between subjective alertness and illuminance has already been established for nocturnal exposures [38]. However, during other times of the day, a dose-response relationship was identified solely based on revised publications, concerning the effect of light on sleepiness [39].

Previous experimental studies on daytime working hours have reported linear relationships between higher illuminance and reduced sleepiness or quicker reaction times [16, 35, 37] and a weak, positive linear relationship between alertness and illuminance levels was found by Smolders, Peeters [40]. Only one experimental study [41] was found that also described an inverted U-shape relationship. This study focused on reaction times in response to melanopic EDI, noting the fastest reaction times at 992 melanopic EDI and slower times at lower and higher melanopic EDI levels. The findings suggest that reaction times could be enhanced at melanopic EDI levels exceeding 595 lx.

The inverted U-shaped relationship observed between melanopic EDI and cognitive performance aligns with models of arousal and performance regulation, such as the Yerkes-Dodson Law [42], which suggests that cognitive performance improves with increasing arousal up to an optimal point, after which excessive stimulation leads to performance deterioration. In the present study, moderate melanopic EDI (70 lx) was associated with the highest accuracy in the n-back task and the lowest perceived workload, suggesting that this level of light optimally engages attentional and cognitive resources without inducing excessive mental strain. At very dim light levels (1 lx and 10 lx), insufficient photic stimulation may fail to elicit the necessary alertness required for cognitive engagement, whereas at very high levels (595 lx), the brightness itself may have been visually disruptive or uncomfortable, increasing mental effort and contributing to performance deterioration. This pattern highlights the complexity of light’s influence on cognition, where neither too little nor excessive brightness may support optimal cognitive performance.

The observed discrepancy between PVT and n-back task performance suggests a speed-accuracy tradeoff, wherein participants may prioritise vigilance and PVT-speed reaction time over working memory performance (n-back) in highly demanding conditions. Under extreme light levels, heightened arousal might facilitate faster but less precise responses, reflecting a shift in cognitive strategy to accommodate the increased mental workload. This finding supports prior research indicating that higher melanopic light levels may enhance simple attentional tasks but do not necessarily improve complex cognitive functions that rely on executive control. Additionally, the type of cognitive demand plays a crucial role in how light modulates performance—sustained attention (PVT) may benefit from heightened arousal, while working memory (n-back) requires a balance between alertness and cognitive control. These findings underscore the importance of tailoring light environments to task-specific cognitive demands, rather than assuming a uniform effect across all cognitive domains.

### Limitations

We consider the following limitations:

#### Wide but limited light exposure range

The current study examined a series of four light exposure levels covering very dim (1 lx melanopic EDI) to very bright (595 lx melanopic EDI) in logarithmic spacing. We did not extend the light exposure levels to more than 595 lx melanopic EDI due to technical limitations, and therefore could not determine the shape of the dose-response curve outside of this range.

#### Limited variation in temporal parameters

The study took place in the evening, adjusted to participants’ habitual bedtime. Whether or not the dose-response behaviour is different in different times of day and/or different circadian phases is not clear. Future work should examine how light exposure during different biological times of day can affect cognitive and mental load outcome parameters.

#### Limited generalisability

It is uncertain whether the inverted U-shaped relationship between illuminance and cognitive performance and mental load would also be found under differing conditions. First, only a laboratory environment was tested, and a recent study [43] comparing the effects of dynamic lighting concepts under both laboratory conditions and field conditions reported that the results were not consistent. Secondly, only the late afternoon and evening were studied, and no comparisons can be made to more typical daytime working hours.

### Future directions

The present study provides an important foundation for understanding the non-linear effects of light on cognition, but further work is needed to generalise these findings and develop predictive models. One key opportunity lies in integrating cognition into computational models of light exposure that already consider circadian and physiological outcomes. Such a framework could incorporate factors such as task complexity, time-of-day effects, and individual variability (e.g., chronotype, light sensitivity) to predict how different lighting conditions impact cognitive performance. Additionally, future studies should examine how light exposure interacts with longer-term neurocognitive adaptations, including potential cumulative effects of daily light exposure patterns on cognitive resilience and fatigue. Expanding this work beyond controlled laboratory settings into real-world environments, such as workplaces, classrooms, or healthcare settings, would also be valuable in translating these insights into practical lighting recommendations.

### Conclusion

The study investigated how log-transformed illuminance at very dim (1 lx melanopic EDI), dim (10 lx melanopic EDI), moderately bright (70 lx melanopic EDI) and very bright (595 lx melanopic EDI) levels affected cognitive performance and mental load, including momentary affect, perceived workload and sleepiness during evening hours, while also considering temporal effects. An inverted U-shaped relationship between increasing melanopic EDI levels and cognitive function, as assessed by the 2-back task, complemented by a U-shaped relationship between melanopic EDI levels and perceived workload along with sleepiness, was discovered. Contrarily, an inverted U-shaped relationship was also found between increasing melanopic EDI levels and reaction times in the Psychomotor Vigilance Task (PVT), with the fastest reactions at very dim (melanopic EDI 1) and very bright (melanopic EDI 595), and the slowest at dim (melanopic EDI 10) and moderately bright (melanopic EDI 70) illuminance levels. While such divergent results as a function of the cognitive demand level of the performance tasks have already been described in previous research, to the best of the authors’ knowledge, this is only the second identification of an inverted U-shape in the relationship between cognitive performance and melanopic EDI levels. This finding indicates that the relationship might be dependent on the time of the day, which in this research were the evening hours of the day. Moreover, a positive linear relationship between exposure duration and arousal states as well as sleepiness across all illuminance levels was found, pointing out the critical role of incorporating temporal factors in the design of building environments.

## Acknowledgements

The authors would like to thank the participants for their contribution.

## Funding

This study was partially supported by a TUM-IAS Hans Fischer Fellowship awarded to SR. and BK.

## Data, code and materials availability

This study’s data, code and materials are available under an open-source (GPL) or open-access license (CC-BY) at https://github.com/tscnlab/ReitmayerEtAl_bioRxiv_2025.

## Supplementary Documents

| Name | Content |
| --- | --- |
| Supplementary Document S1 | ▪ Rendering of the TUM SenseLab with the experimental setup for two participants (Figure S1)<br>▪ Pairwise correlations between performance, sleepiness, and workload measures (Figures S2-S8)<br>▪ Correlation matrix (Figure S9)<br>▪ Correlations between the operative temperature (°C) and both performance and sleepiness (Figures S10- S12)<br>▪ Operative temperature and ambient lux levels per participant and session (Figure S13) |
| Supplementary Document S2 | Individual participant analysis showing the relationship between each parameter (KSS, MAS, NASA-TLX, PVT, N-back) and the different lighting scenarios |
| Supplementary Document S3 | Outputs linear mixed model analysis |
| Supplementary Document S4 | ENLIGHT Checklist |

## Author contributions

Conceptualisation: AR, BK, MK, CRL, MS

Methodology: AR, BK, MK, CRL, MS

Software: AR, MS

Validation: AR, BK, MK, CRL, MS

Formal analysis: MS

Investigation: AR, BK, MK, CRL

Resources: TA, MS

Data Curation: AR, BK, MK

Writing - Original Draft: AR, MS

Writing - Review & Editing: AR, BK, KJ, CM, MMC, TA, SR, MS

Visualisation: AR

Supervision: KJ, CM, MMC, TA, SR, MS

Project administration: BK, MS

Funding acquisition: BK, TA, SR, MS

## Notes

https://github.com/tscnlab/ReitmayerEtAl_bioRxiv_2025

### Competing Interest Statement

The authors have declared no competing interest.

## References

1. Blume, C., C. Garbazza, and M. Spitschan, Effects of light on human circadian rhythms, sleep and mood. Somnologie (Berl), 2019. 23(3): p. 147–156.

2. Brown, T.M., et al., Recommendations for daytime, evening, and nighttime indoor light exposure to best support physiology, sleep, and wakefulness in healthy adults. PLoS Biol, 2022. 20(3): p. e3001571.

3. Vetter, C., et al., A Review of Human Physiological Responses to Light: Implications for the Development of Integrative Lighting Solutions. Leukos, 2021. 18(3): p. 387–414.

4. Brown, T.M., Melanopic illuminance defines the magnitude of human circadian light responses under a wide range of conditions. J Pineal Res, 2020. 69(1): p. e12655.

5. Cajochen, C., Alerting effects of light. Sleep Med Rev, 2007. 11(6): p. 453–64.

6. Cajochen, C., et al., Dose-response relationship for light intensity and ocular and electroencephalographic correlates of human alertness. Behav Brain Res, 2000. 115(1): p. 75–83.

7. Lok, R., et al., Light, Alertness, and Alerting Effects of White Light: A Literature Overview. J Biol Rhythms, 2018. 33(6): p. 589–601.

8. Lucas, R.J., et al., Measuring and using light in the melanopsin age. Trends Neurosci, 2014. 37(1): p. 1–9.

9. Spitschan, M., Melanopsin contributions to non-visual and visual function. Curr Opin Behav Sci, 2019. 30: p. 67–72.

10. Mahoney, H.L. and T.M. Schmidt, The cognitive impact of light: illuminating ipRGC circuit mechanisms. Nat Rev Neurosci, 2024. 25(3): p. 159–175.

11. Huiberts, L.M., K.C.H.J. Smolders, and Y.A.W. de Kort, Non-image forming effects of illuminance level: Exploring parallel effects on physiological arousal and task performance. Physiology & Behavior, 2016. 164: p. 129–139.

12. Lok, R., et al., Light, Alertness, and Alerting Effects of White Light: A Literature Overview. Journal of Biological Rhythms, 2018. 33(6): p. 589–601.

13. Souman, J.L., et al., Acute alerting effects of light: A systematic literature review. Behavioural Brain Research, 2018. 337: p. 228–239.

14. Houser, K.W. and T. Esposito, Human-Centric Lighting: Foundational Considerations and a Five-Step Design Process. Front Neurol, 2021. 12: p. 630553.

15. Dinges, D.F. and J.W. Powell, Microcomputer analyses of performance on a portable, simple visual RT task during sustained operations. Behavior Research Methods, Instruments, & Computers, 1985. 17(6): p. 652–655.

16. Phipps-Nelson, J., et al., Daytime Exposure to Bright Light, as Compared to Dim Light, Decreases Sleepiness and Improves Psychomotor Vigilance Performance. Sleep, 2003. 26(6): p. 695–700.

17. Badia, P., et al., Bright light effects on body temperature, alertness, EEG and behavior. Physiology & Behavior, 1991. 50(3): p. 583–588.

18. Zhu, Y., et al., Effects of Illuminance and Correlated Color Temperature on Daytime Cognitive Performance, Subjective Mood, and Alertness in Healthy Adults. Environment and Behavior, 2017. 51(2): p. 199–230.

19. French, J., P. Hannon, and G.C. Brainard, Effects of bright illuminance on body temperature and human performance. Annual review of chronopharmacology, 1990. 7.

20. Zhou, A. and Y. Pan, Effects of indoor lighting environments on paper reading efficiency and brain fatigue: an experimental study. Frontiers in Built Environment, 2023. 9.

21. Spitschan, M., S. Nam, and J.A. Veitch. luox: Platform for calculating quantities related to light and lighting [Software]. 2022; Available from: https://luox.app/.

22. Spitschan, M., et al., ENLIGHT: A consensus checklist for reporting laboratory-based studies on the non-visual effects of light in humans. EBioMedicine, 2023. 98: p. 104889.

23. ANSI/ASHRAE, Standard 55: 2023, Thermal Environmental Conditions for Human Occupancy. 2023, Atlanta, USA: American National Standards Institute (ANSI), American National Standards Institute.

24. Owen, A.M., et al., N-back working memory paradigm: a meta-analysis of normative functional neuroimaging studies. Hum Brain Mapp, 2005. 25(1): p. 46–59.

25. Stoet, G., PsyToolkit: A software package for programming psychological experiments using Linux. Behavior Research Methods, 2010. 42(4): p. 1096–1104.

26. Stoet, G., PsyToolkit: A Novel Web-Based Method for Running Online Questionnaires and Reaction-Time Experiments. Teaching of Psychology, 2016. 44(1): p. 24–31.

27. Jaeggi, S.M., et al., The concurrent validity of the N-back task as a working memory measure. Memory, 2010. 18(4): p. 394–412.

28. Dorrian, J., N. Rogers, and D. Dinges, Psychomotor vigilance perfomance: neurocognitive assay sensitive to sleep loss In: Kushida C, ed. Sleep deprivation: clinical issues, pharmacology and sleep loss effects. 2005, New York, Marcel Dekker. p. pp. 39–70.

29. Van Dongen, H.P.A., et al., The Cumulative Cost of Additional Wakefulness: Dose-Response Effects on Neurobehavioral Functions and Sleep Physiology From Chronic Sleep Restriction and Total Sleep Deprivation. Sleep, 2003. 26(2): p. 117–126.

30. Hart, S.G. and L.E. Staveland, Development of NASA-TLX (Task Load Index): Results of Empirical and Theoretical Research, in Advances in Psychology, P.A. Hancock and N. Meshkati, Editors. 1988, North-Holland. p. 139–183.

31. Grier, R.A., How High is High? A Meta-Analysis of NASA-TLX Global Workload Scores. Proceedings of the Human Factors and Ergonomics Society Annual Meeting, 2015. 59(1): p. 1727–1731.

32. Gee, P., et al., Chapter 5 Measuring Affect Over Time: The Momentary Affect Scale, in Experiencing and Managing Emotions in the Workplace, N.M. Ashkanasy, C.E.J. Härtel, and W.J. Zerbe, Editors. 2012, Emerald Group Publishing Limited. p. 141–173.

33. Kaida, K., et al., Validation of the Karolinska sleepiness scale against performance and EEG variables. Clinical Neurophysiology, 2006. 117(7): p. 1574–1581.

34. Team, R.C., R: A language and environment for statistical computing. 2023, R Foundation for Statistical Computing: Vienna, Austria.

35. Smolders, K.C.H.J., Y.A.W. de Kort, and P.J.M. Cluitmans, A higher illuminance induces alertness even during office hours: Findings on subjective measures, task performance and heart rate measures. Physiology & Behavior, 2012. 107(1): p. 7–16.

36. Huiberts, L.M., K.C.H.J. Smolders, and Y.A.W. de Kort, Shining light on memory: Effects of bright light on working memory performance. Behavioural Brain Research, 2015. 294: p. 234–245.

37. Smolders, K.C.H.J. and Y.A.W. de Kort, Bright light and mental fatigue: Effects on alertness, vitality, performance and physiological arousal. Journal of Environmental Psychology, 2014. 39: p. 77–91.

38. Cajochen, C., et al., Dose-response relationship for light intensity and ocular and electroencephalographic correlates of human alertness. Behavioural Brain Research, 2000. 115(1): p. 75–83.

39. Hommes, V. and M.C. Giménez, A revision of existing Karolinska Sleepiness Scale responses to light: A melanopic perspective. Chronobiology International, 2015. 32(6): p. 750–756.

40. Smolders, K.C.H.J., et al., Investigation of Dose-Response Relationships for Effects of White Light Exposure on Correlates of Alertness and Executive Control during Regular Daytime Working Hours. Journal of Biological Rhythms, 2018. 33(6): p. 649–661.

41. Münch, M., et al. Non-Linear Relationship for Reaction Time and Melanopic EDI—Is There an Optimum for Office Lighting? in 34th Annual Meeting of the Society for Light Treatment and Biological Rhythms (SLTBR). 2023. Lausanne, Switzerland: Clocks Sleep.

42. Yerkes, R.M. and J.D. Dodson, The relation of strength of stimulus to rapidity of habit-formation. Journal of Comparative Neurology and Psychology, 2004. 18(5): p. 459–482.

43. Aries, M.B.C., F. Beute, and G. Fischl, Assessment protocol and effects of two dynamic light patterns on human well-being and performance in a simulated and operational office environment. Journal of Environmental Psychology, 2020. 69: p. 101409.

