## Supplementary Document S1 for "Non-linear effects of evening light exposure on cognitive performance"

Document S1 Figures S1-13

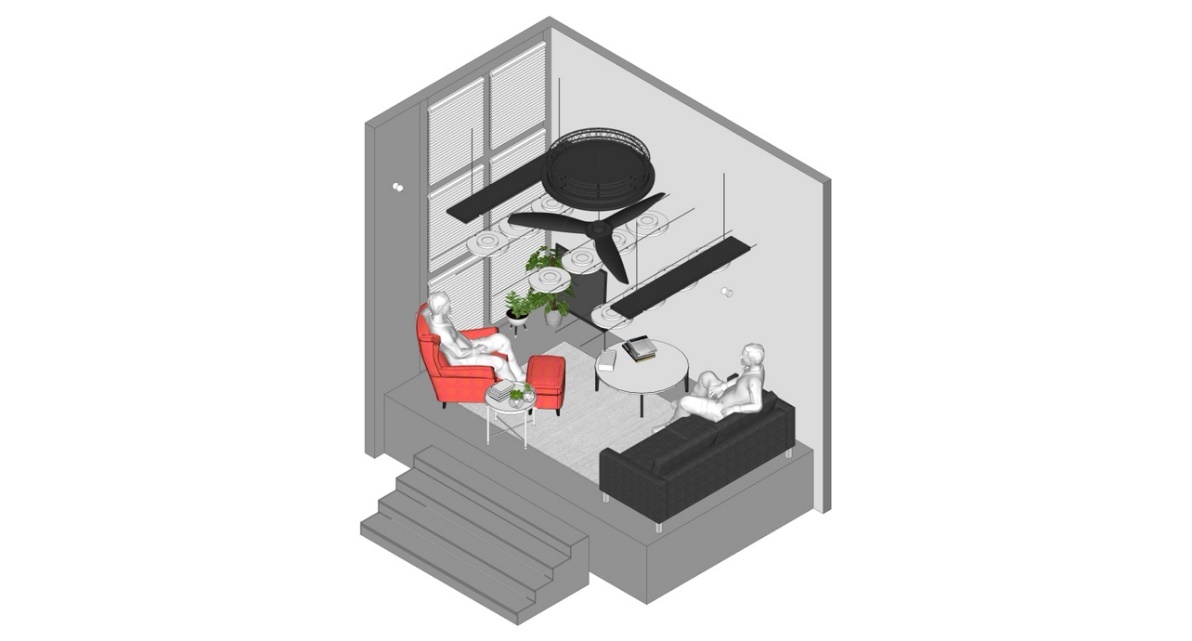

Figure S1 Rendering of the TUM SenseLab with the experimental setup for two participants

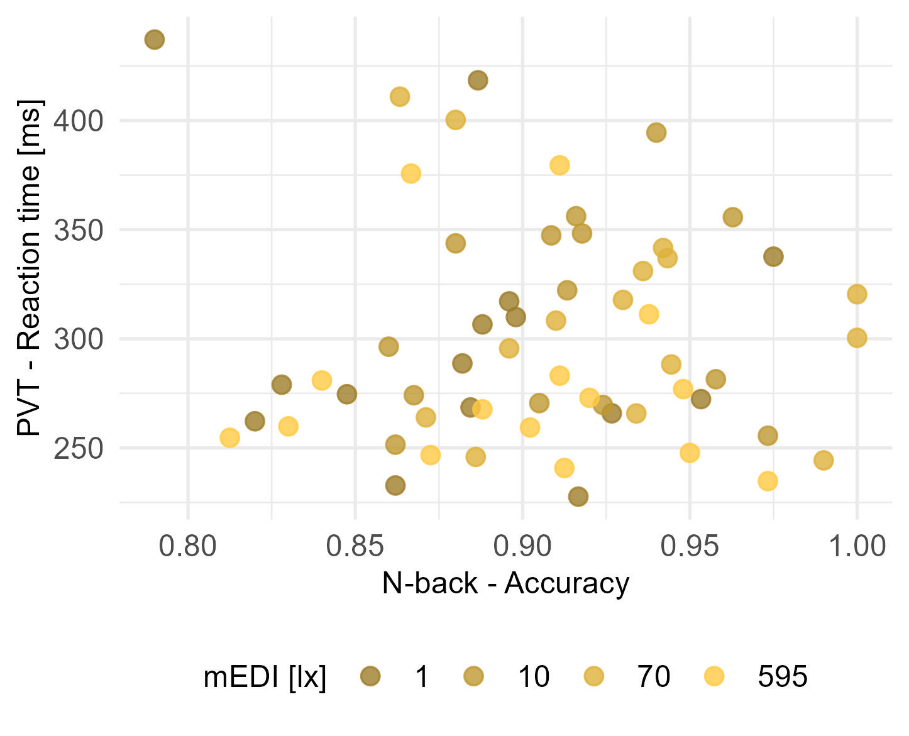

Figure S2 Correlation between reaction time (PVT) and accuracy (n-back)

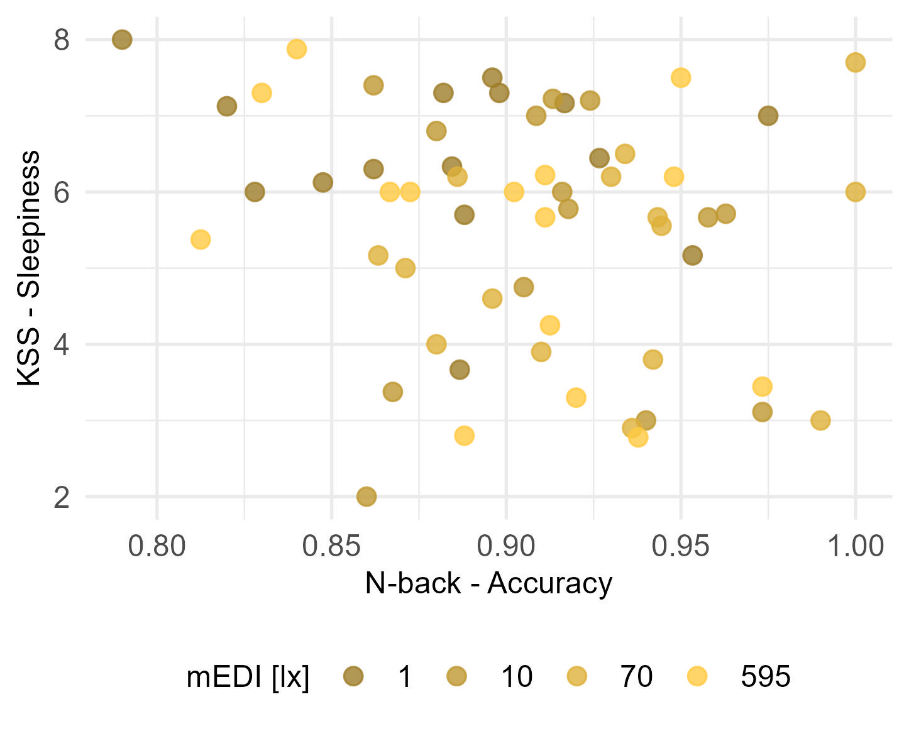

Figure S3 Correlation between sleepiness (KSS) and accuracy (n-back)

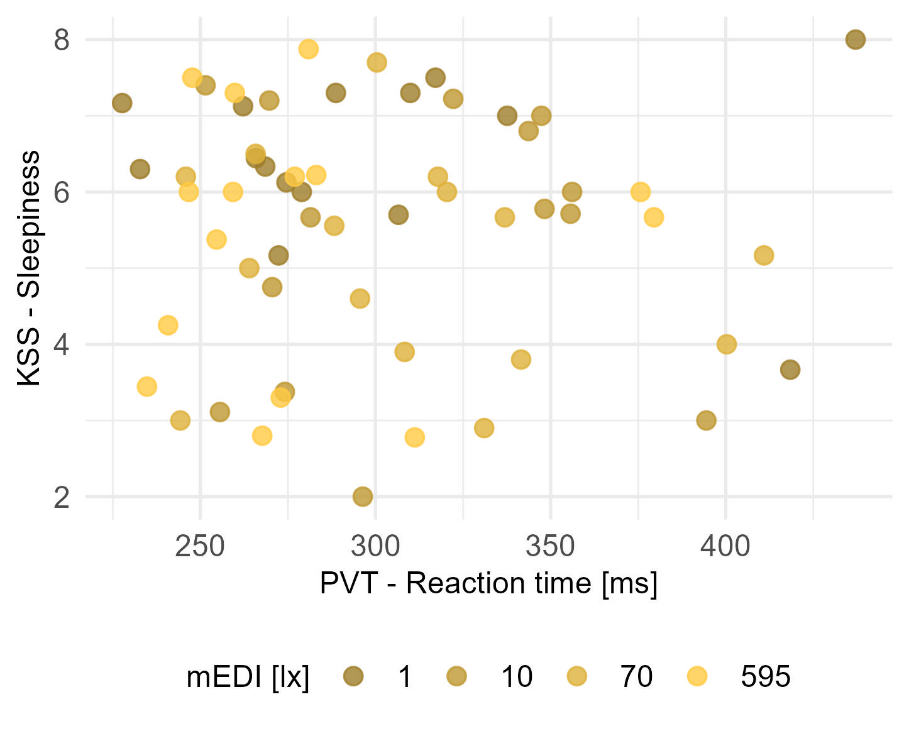

Figure S4 Correlation between sleepiness (KSS) and reaction time (PVT)

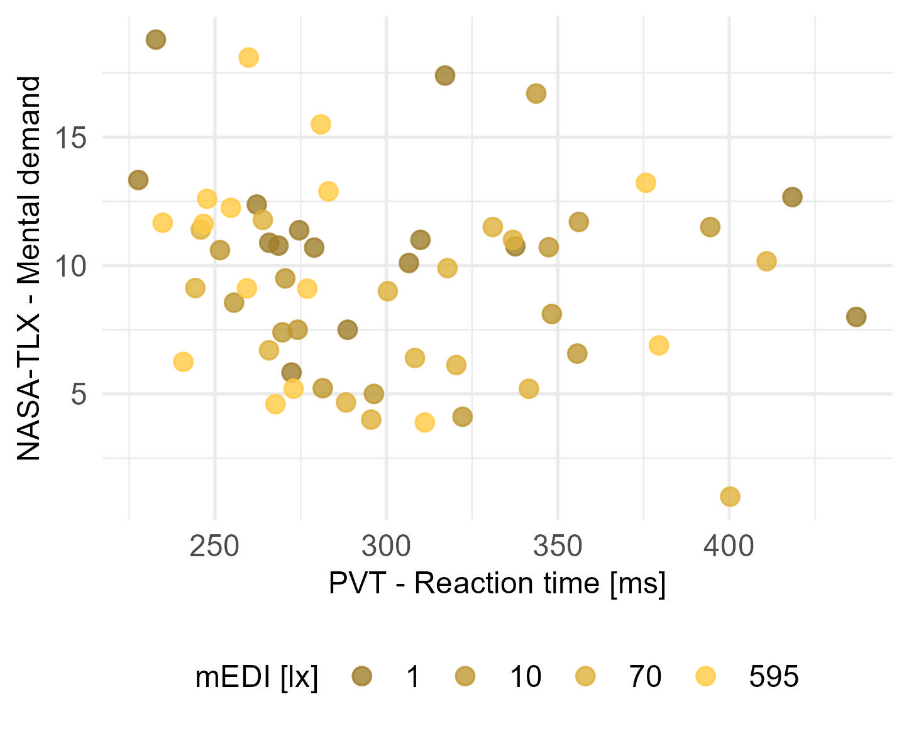

Figure S5 Correlation between mental demand (NASA-TLX) and reaction time (PVT)

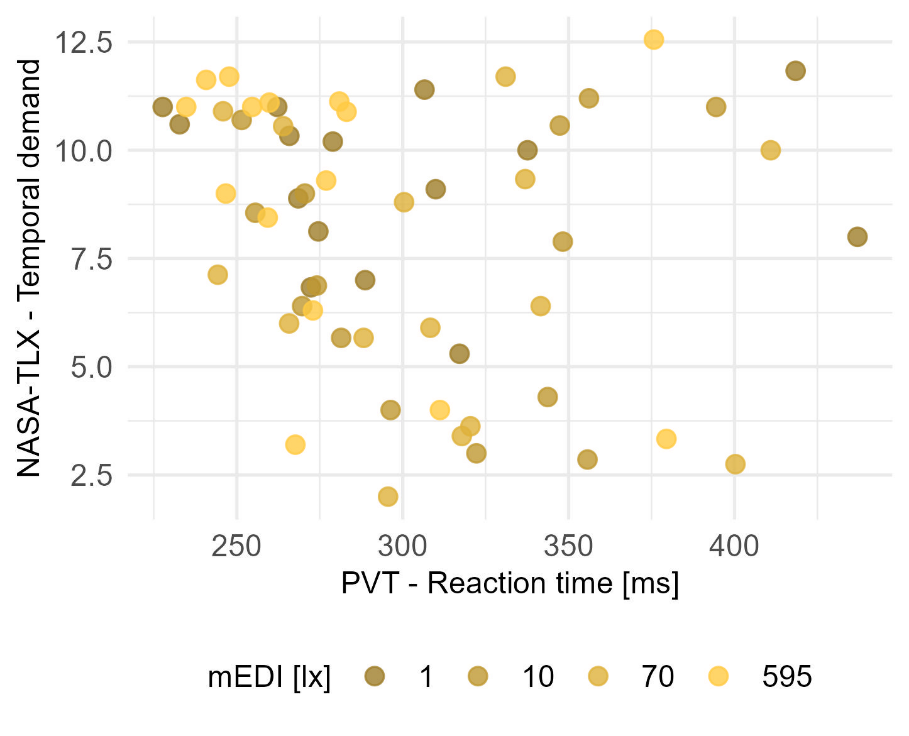

Figure S6 Correlation between temporal demand (NASA-TLX) and reaction time (PVT)

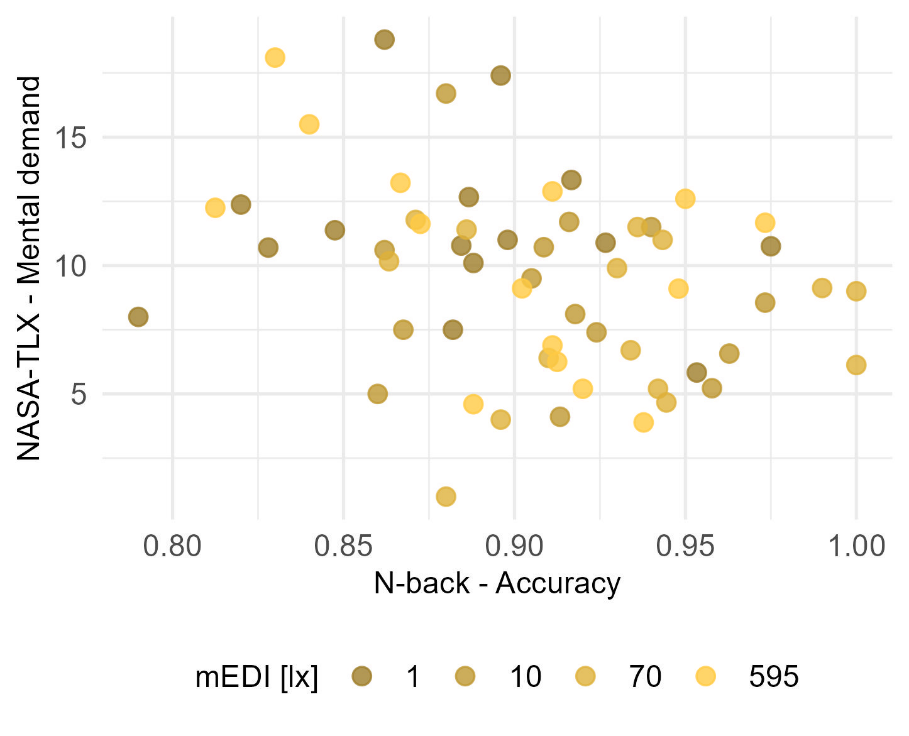

Figure S7 Correlation between mental demand (NASA-TLX) and accuracy (n-back)

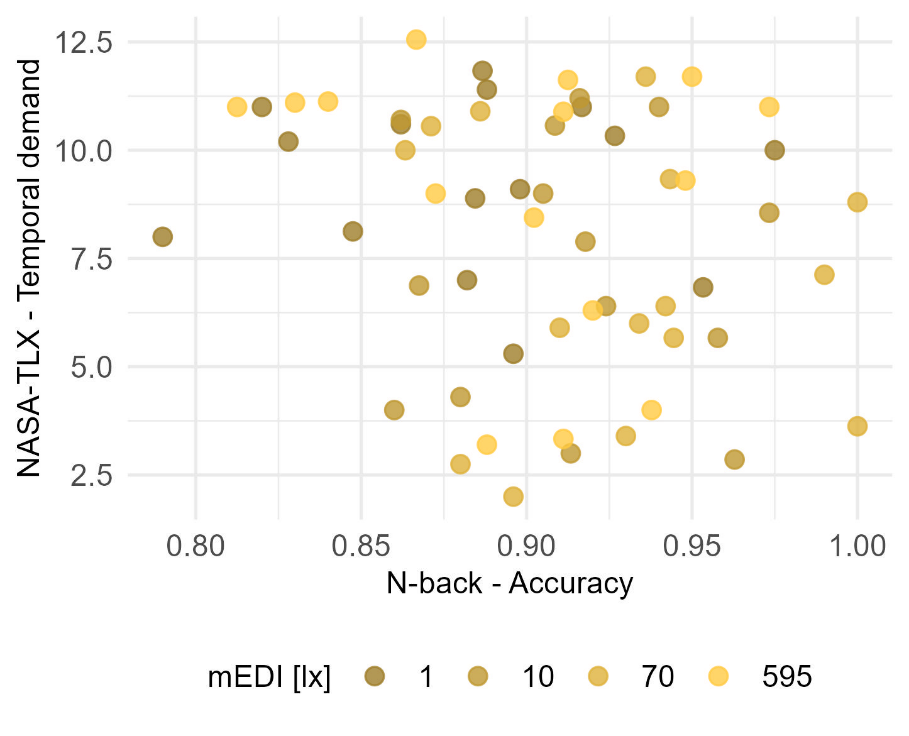

Figure S8 Correlation between temporal demand (NASA-TLX) and accuracy (n-back)

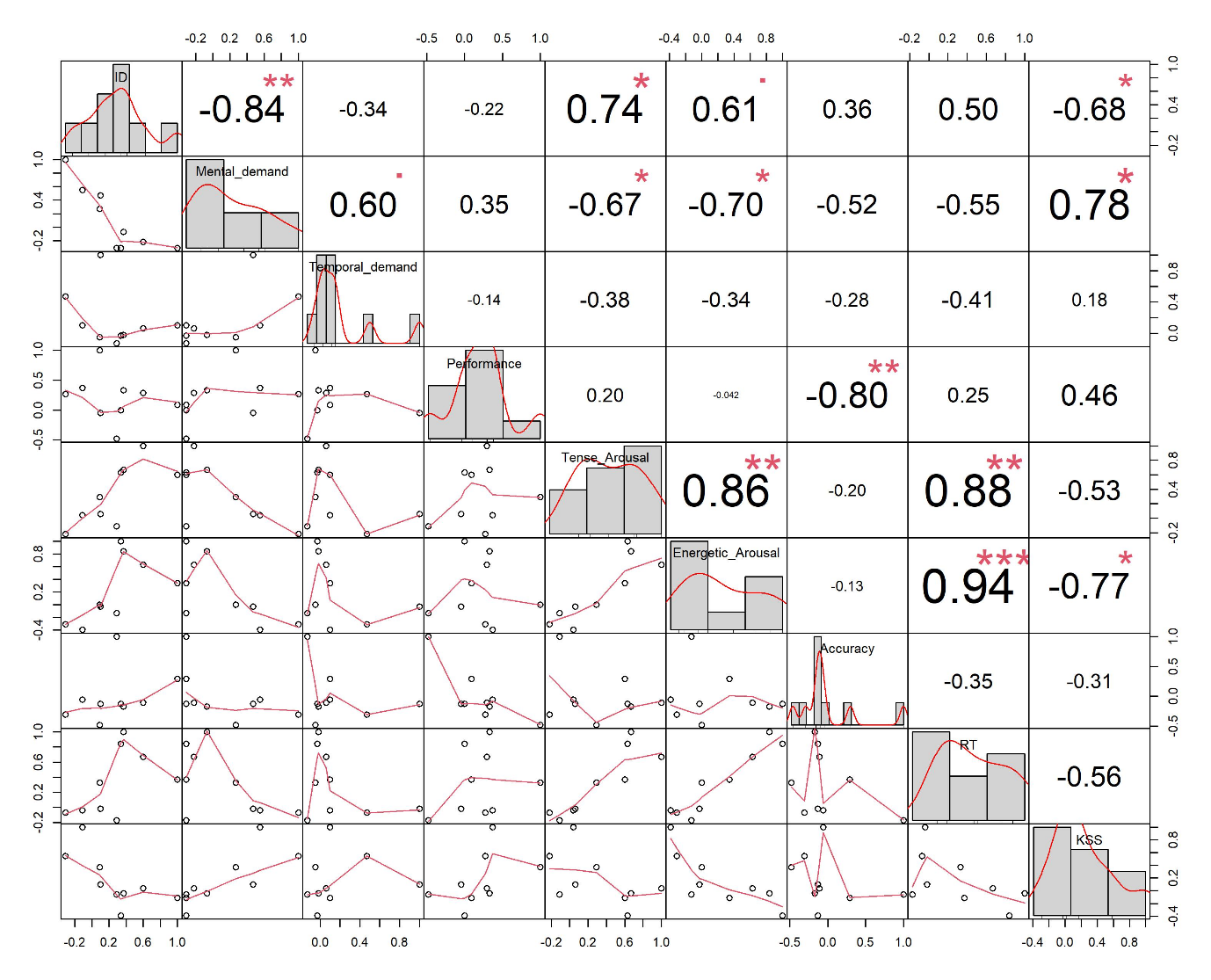

Figure S9 Correlation matrix

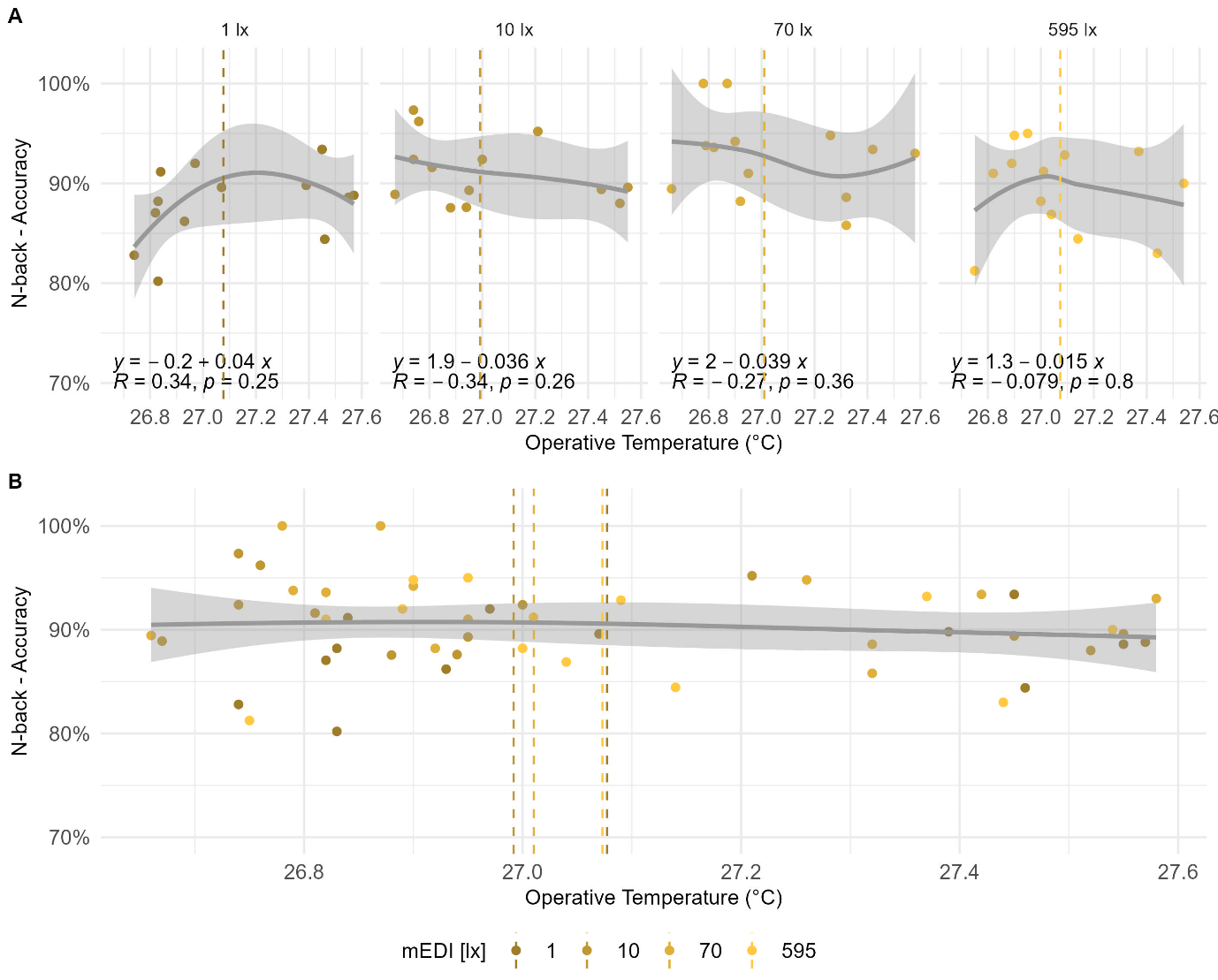

Figure S10 Correlations between the operative temperature (°C) and n-back task performance (A) categorised according to the four scenarios and (B) combined data

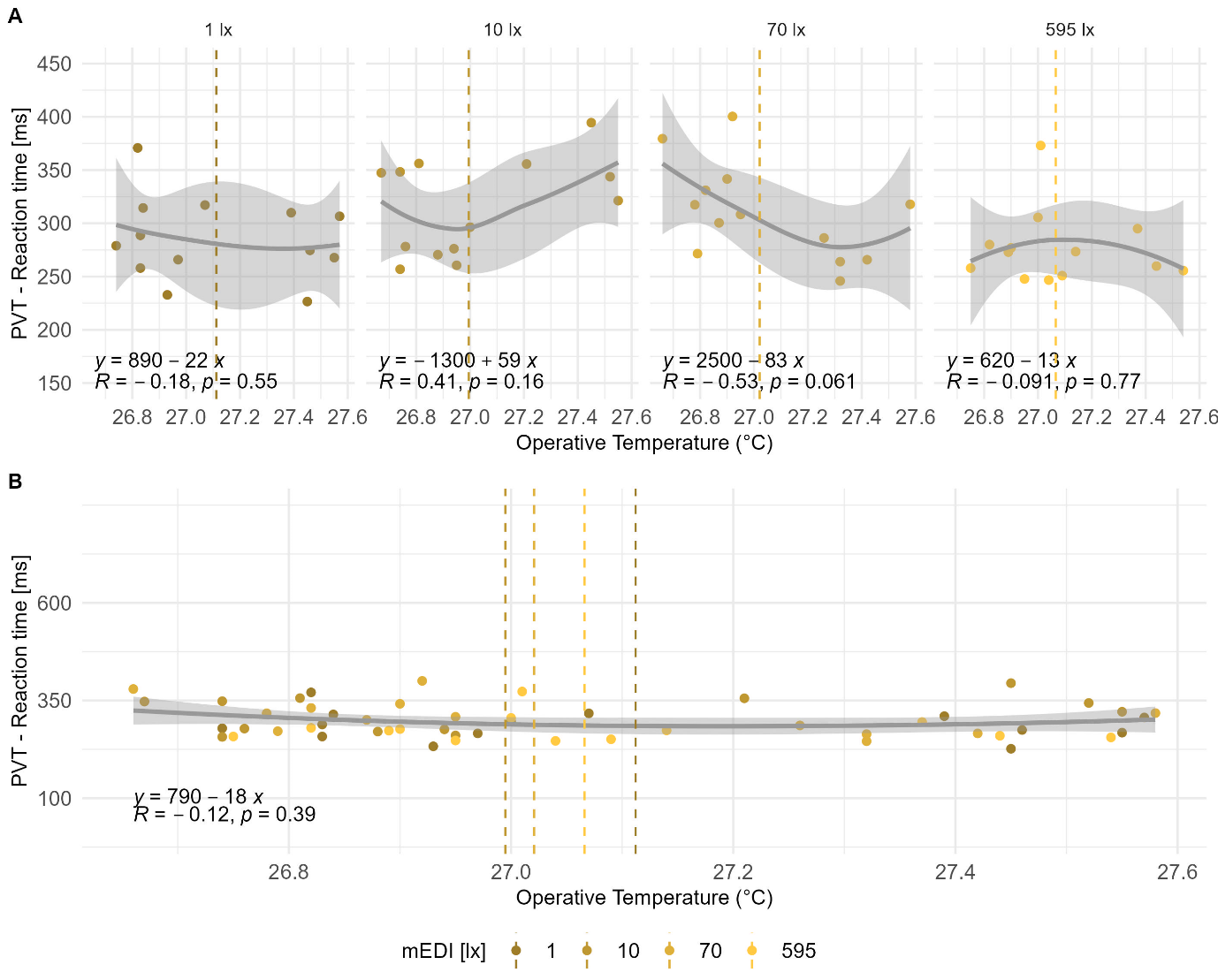

Figure S11 Correlations between the operative temperature (°C) and PVT reaction times (A) categorised according to the four scenarios and (B) combined data.

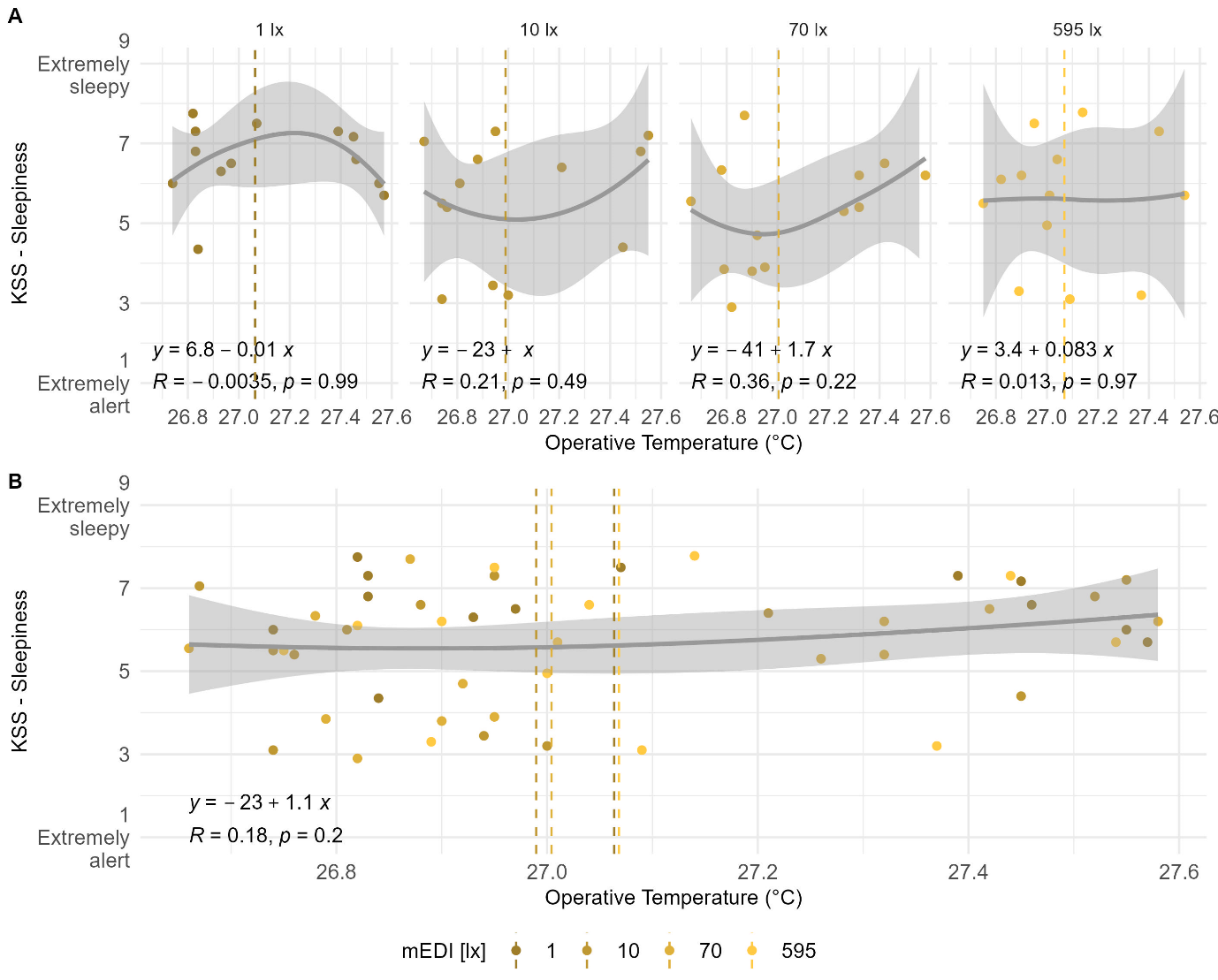

Figure S12 Correlations between the operative temperature (°C) and KSS sleepiness (A) categorised according to the four scenarios and (B) combined data.

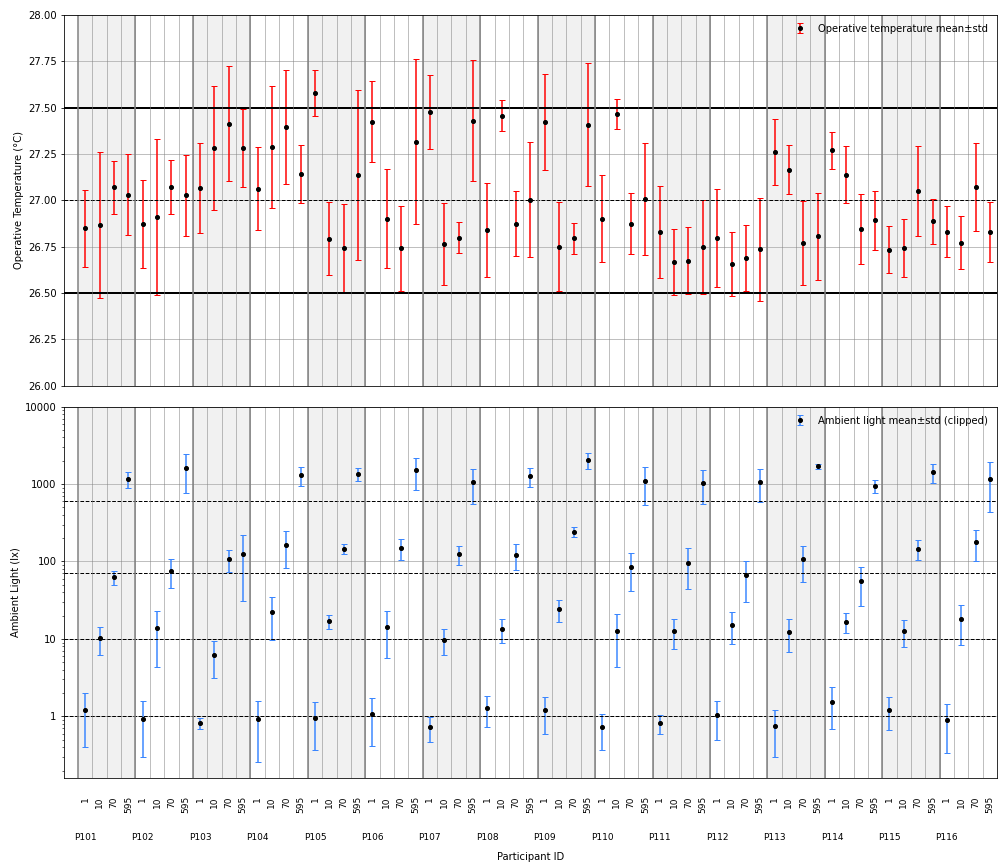

Figure S13 Operative temperature and ambient lux levels per participant and session.
