## Supplementary Document S2 for "Non-linear effects of evening light exposure on cognitive performance"

Document S2 Matrix

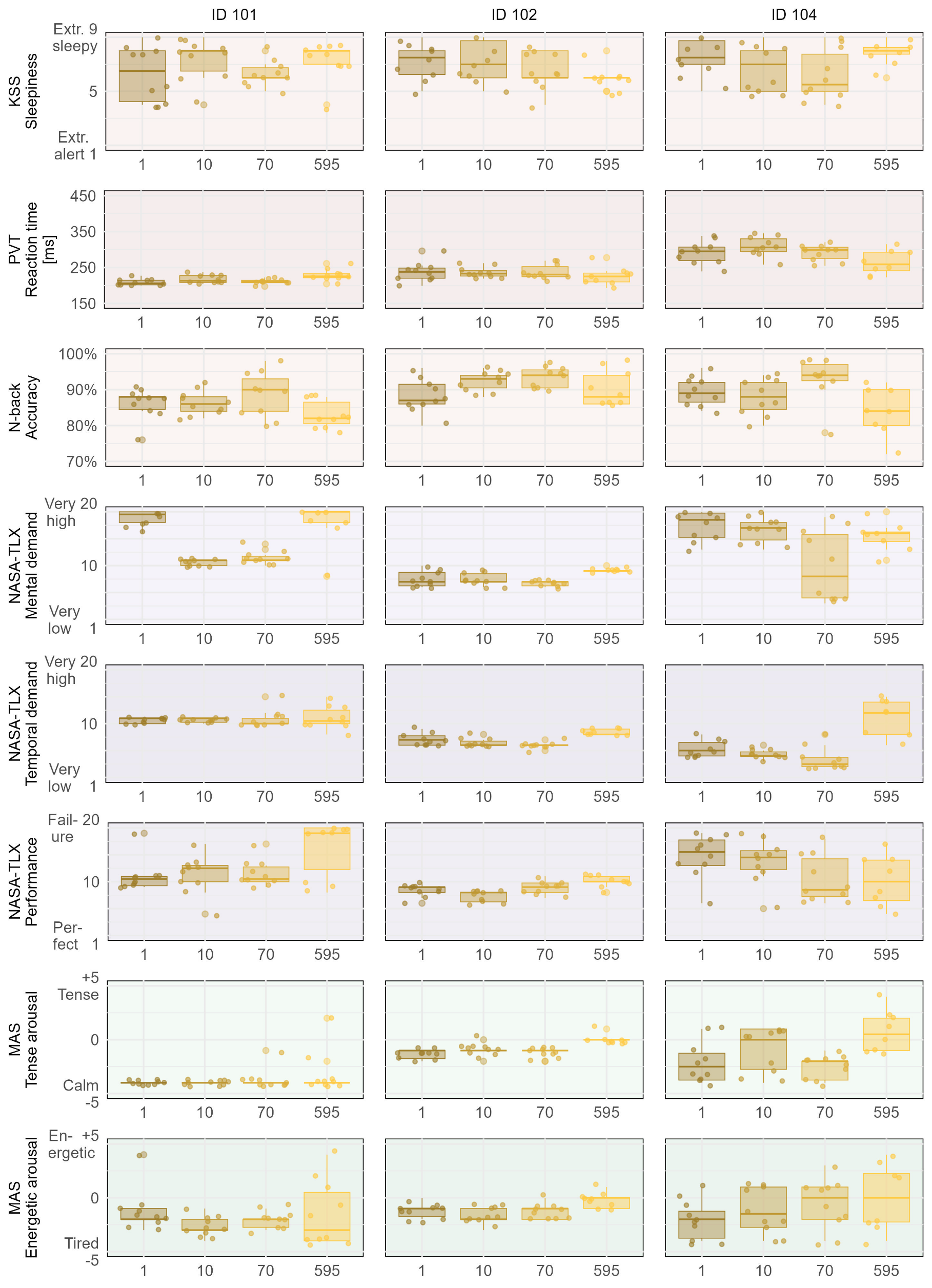

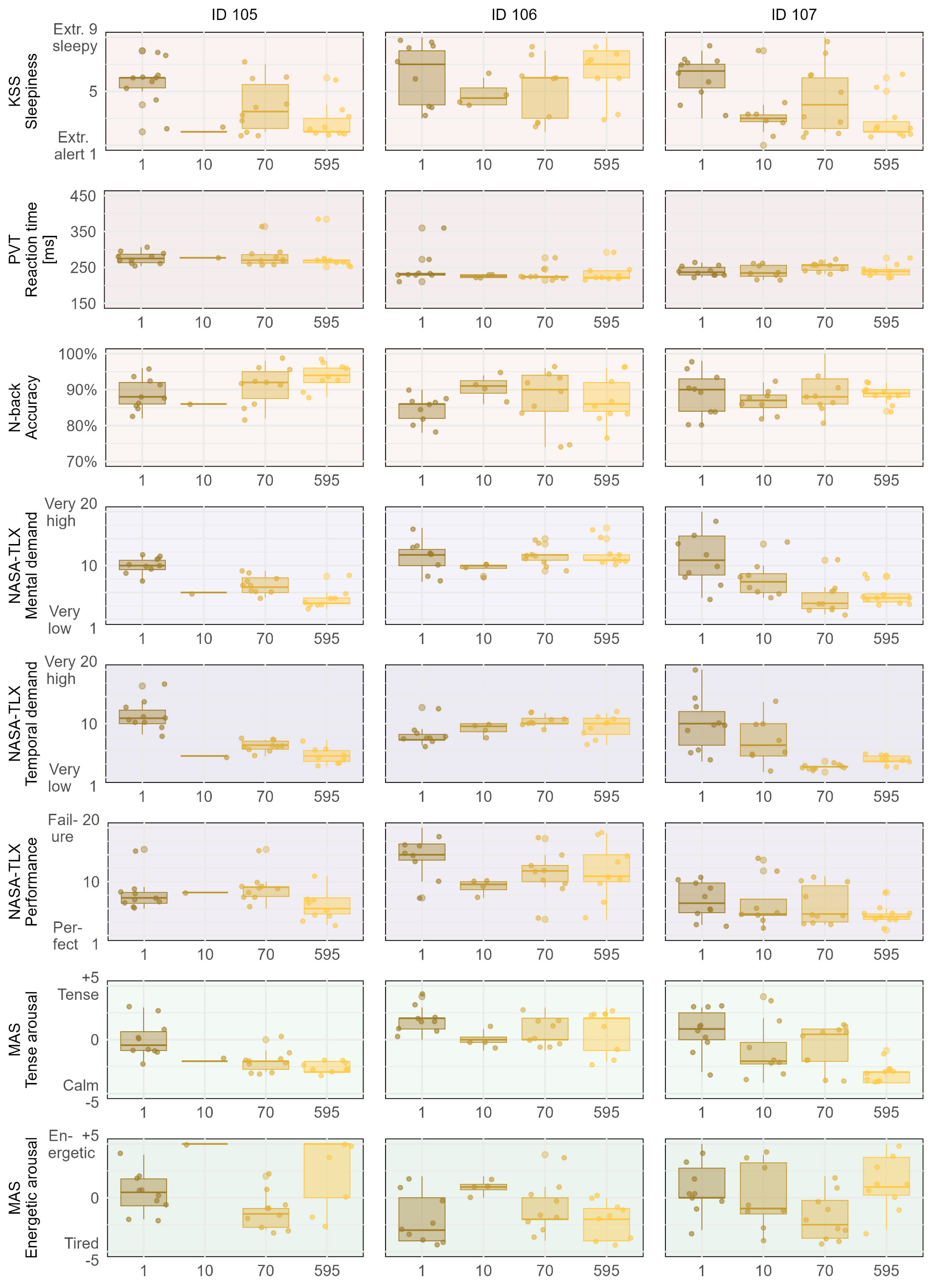

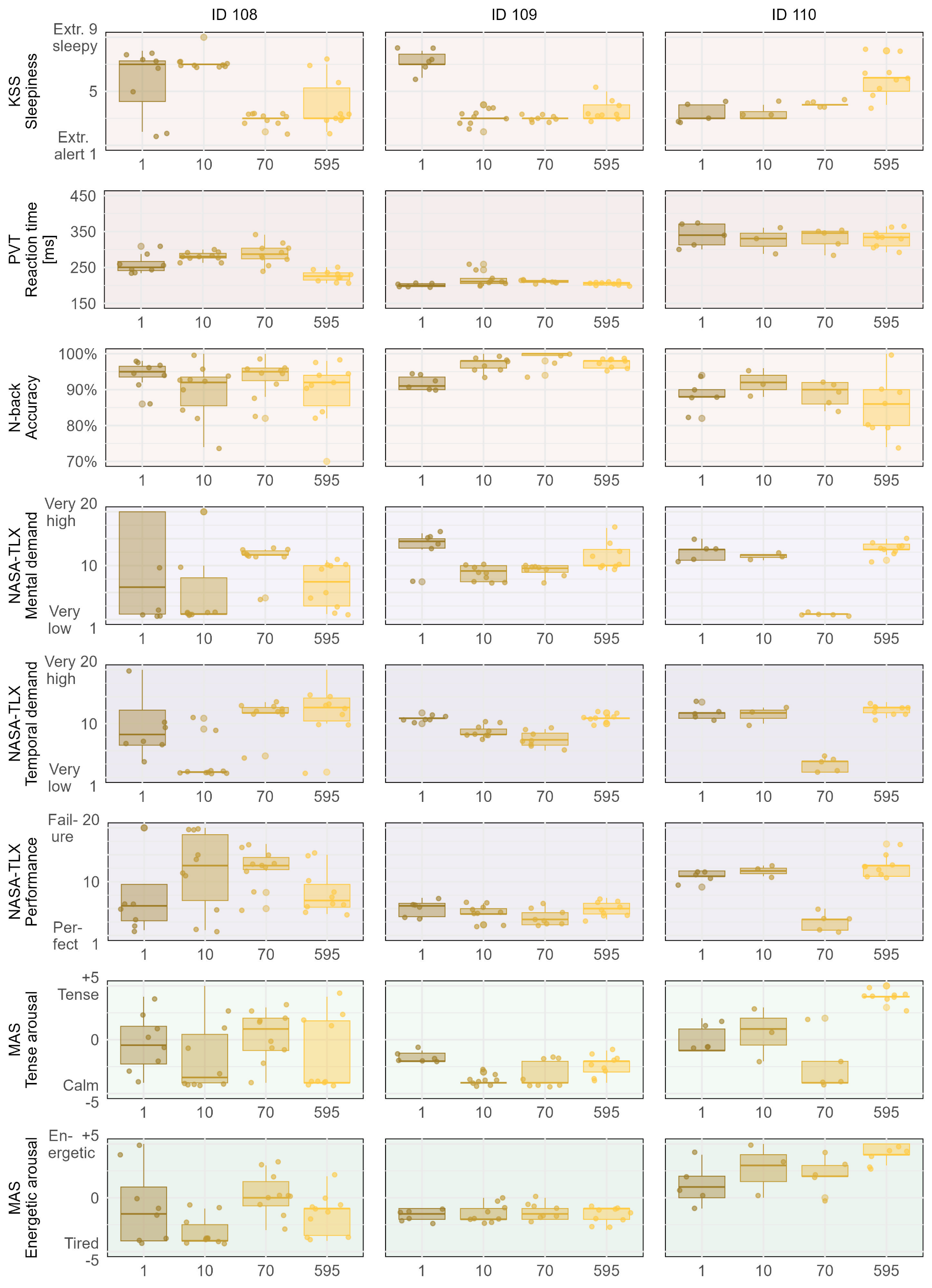

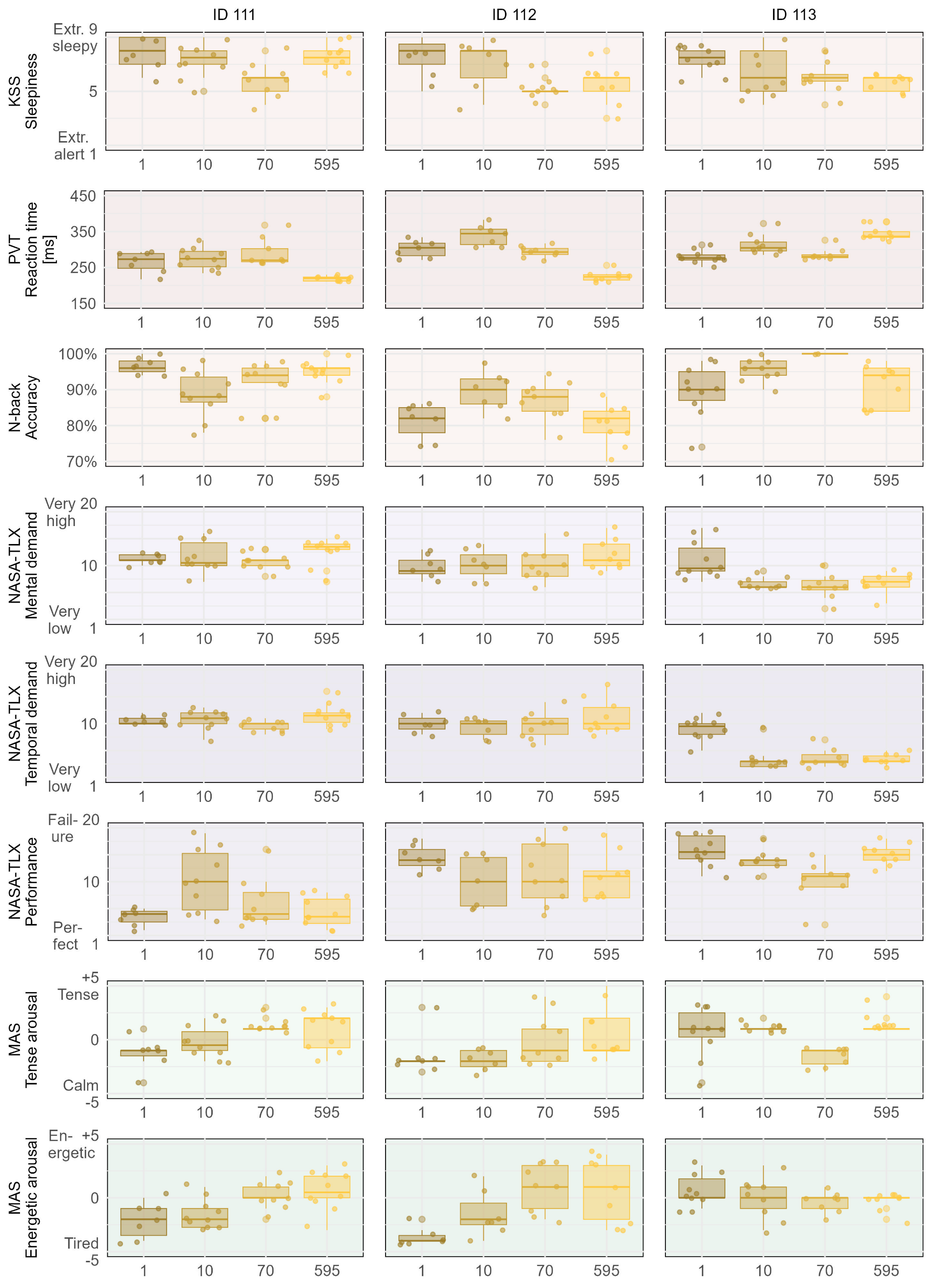

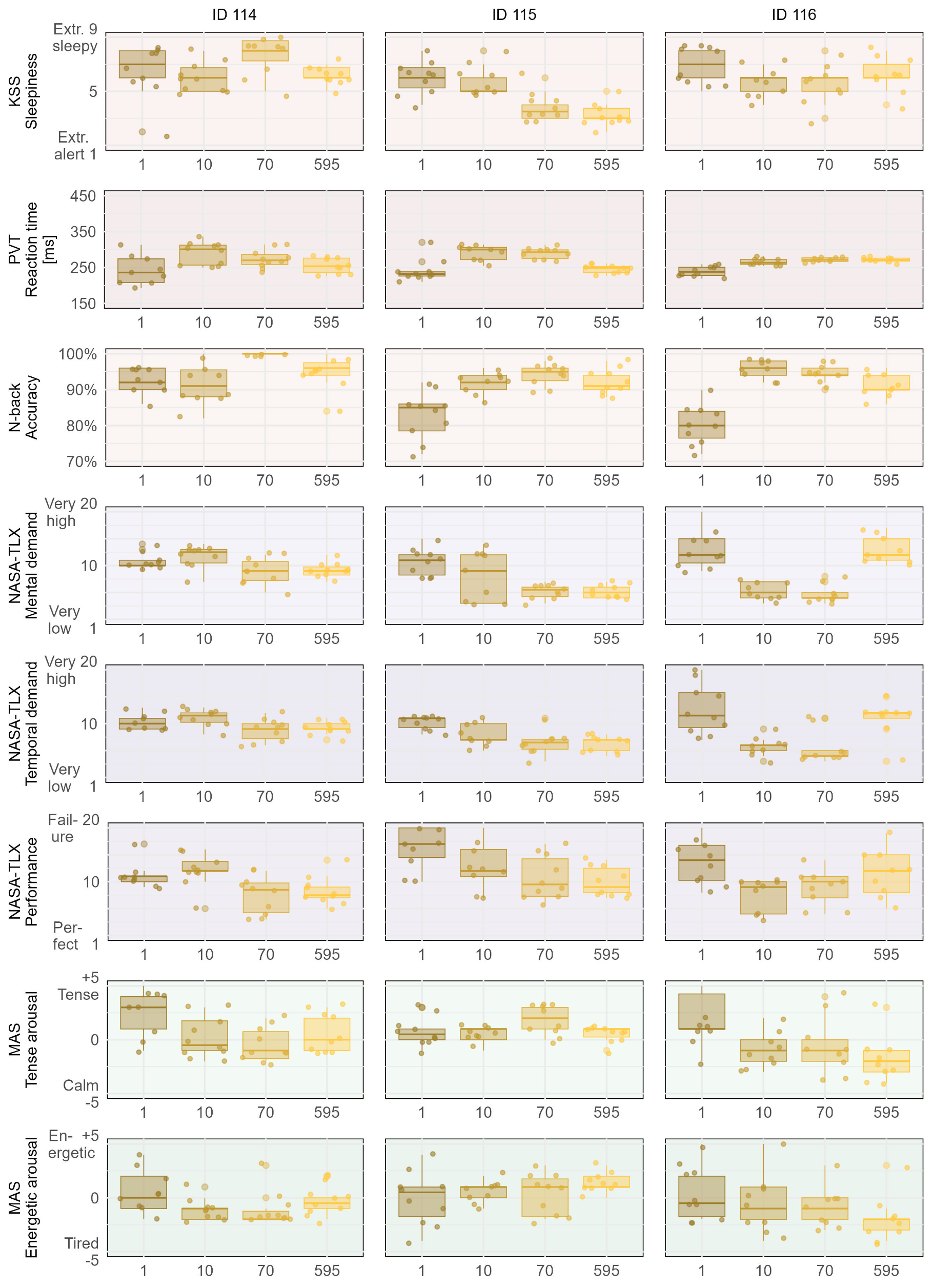
