## Supplementary Document S3 for "Non-linear effects of evening light exposure on cognitive performance"

Document S3 Table S1

Table S1 Outputs linear mixed model analysis

| **PVT Reaction time** |  |  |  | |  | |  | |  |
| --- | --- | --- | --- | --- | --- | --- | --- | --- | --- |
| LMM | Estimate | Std. Error | df | t value | | Pr(>\|t\|) | | Bonferroni adj. p | |
| (Intercept) | 262 | 12.5 | 28.8 | | 20.8 | | < 0.00 | | < 0.00 |
| Scenario | 46.9 | 7.12 | 502 | | 6.58 | | < 0.00 | | < 0.00 |
| I(Scenario^2) | -17.7 | 2.10 | 502 | | -8.44 | | < 0.00 | | < 0.00 |
| Timepoint | 4.93 | 1.09 | 502 | | 4.52 | | < 0.00 | | < 0.00 |
| Scenario:Timepoint | -0.32 | 0.603 | 502 | | -0.531 | | 0.596 | | 1 |
| **KSS Sleepiness** |  |  |  | |  | |  | |  |
|  | Estimate | Std. Error | df | | t value | | Pr(>\|t\|) | | Bonferroni adj. p |
| (Intercept) | 5.45 | 0.380 | 35.4 | | 14.3 | | < 0.00 | | < 0.00 |
| Scenario | -1.20 | 0.245 | 574 | | -4.92 | | < 0.00 | | < 0.00 |
| I(Scenario^2) | 0.249 | 0.073 | 574 | | 3.38 | | < 0.00 | | 0.003 |
| Timepoint | 0.195 | 0.037 | 574 | | 5.21 | | < 0.00 | | < 0.00 |
| Scenario:Timepoint | 0.016 | 0.021 | 574 | | 0.744 | | 0.457 | | 1 |
| **MAS Tense arousal** | |  |  | |  | |  | |  |
|  | Estimate | Std. Error | df | | t value | | Pr(>\|t\|) | | Bonferroni adj. p |
| (Intercept) | 5.20 | 0.465 | 40.5 | | 11.1 | | < 0.00 | | < 0.00 |
| Scenario | -1.06 | 0.315 | 573 | | -3.36 | | < 0.00 | | 0.004 |
| I(Scenario^2) | 0.369 | 0.094 | 573 | | 3.91 | | < 0.00 | | < 0.00 |
| Timepoint | -0.084 | 0.047 | 573 | | -1.77 | | 0.076 | | 0.382 |
| Scenario:Timepoint | 0.005 | 0.027 | 573 | | 0.190 | | 0.849 | | 1 |
| **MAS Energetic arousal** | |  |  | |  | |  | |  |
|  | Estimate | Std. Error | df | | t value | | Pr(>\|t\|) | | Bonferroni adj. p |
| (Intercept) | 6.08 | 0.400 | 68.2 | | 15.1 | | < 0.00 | | < 0.00 |
| Scenario | -0.440 | 0.310 | 567 | | -1.41 | | 0.157 | | 0.786 |
| I(Scenario^2) | 0.240 | 0.092 | 567 | | 2.59 | | 0.009 | | 0.047 |
| Timepoint | -0.322 | 0.047 | 567 | | -6.85 | | < 0.00 | | < 0.00 |
| Scenario:Timepoint | 0.008 | 0.027 | 567 | | 0.320 | | 0.748 | | 1 |
| **NASA-TLX Mental demand** | |  |  | |  | |  | |  |
|  | Estimate | Std. Error | df | | t value | | Pr(>\|t\|) | | Bonferroni adj. p |
| (Intercept) | 12.3 | 0.874 | 36.7 | | 14.0 | | < 0.00 | | < 0.00 |
| Scenario | -5.01 | 0.572 | 576 | | -8.76 | | < 0.00 | | < 0.00 |
| I(Scenario^2) | 1.46 | 0.171 | 576 | | 8.55 | | < 0.00 | | < 0.00 |
| Timepoint | -0.013 | 0.086 | 576 | | -0.161 | | 0.872 | | 1 |
| Scenario:Timepoint | 0.027 | 0.050 | 576 | | 0.555 | | 0.579 | | 1 |
| **NASA-TLX Temporal demand** | |  |  | |  | |  | |  |
|  | Estimate | Std. Error | df | | t value | | Pr(>\|t\|) | | Bonferroni adj. p |
| (Intercept) | 10.8 | 0.682 | 40.4 | | 15.8 | | < 0.00 | | < 0.00 |
| Scenario | -3.93 | 0.455 | 570 | | -8.62 | | < 0.00 | | < 0.00 |
| I(Scenario^2) | 1.12 | 0.135 | 570 | | 8.32 | | < 0.00 | | < 0.00 |
| Timepoint | -0.189 | 0.069 | 570 | | -2.72 | | 0.006 | | 0.033 |
| Scenario:Timepoint | 0.087 | 0.040 | 570 | | 2.19 | | 0.028 | | 0.143 |
| **NASA-TLX Performance** | |  |  | |  | |  | |  |
|  | Estimate | Std. Error | df | | t value | | Pr(>\|t\|) | | Bonferroni adj. p |
| (Intercept) | 10.9 | 0.969 | 41.1 | | 11.3 | | < 0.00 | | < 0.00 |
| Scenario | -2.12 | 0.661 | 576 | | -3.20 | | 0.001 | | 0.007 |
| I(Scenario^2) | 0.476 | 0.198 | 576 | | 2.40 | | 0.016 | | 0.082 |
| Timepoint | 0.025 | 0.100 | 576 | | 0.253 | | 0.800 | | 1 |
| Scenario:Timepoint | 0.025 | 0.057 | 576 | | 0.448 | | 0.654 | | 1 |
| **N-back Accuracy** |  |  |  | |  | |  | |  |
|  | Estimate | Std. Error | df | | t value | | Pr(>\|t\|) | | Bonferroni adj. p |
| (Intercept) | 0.864 | 0.012 | 49.3 | | 71.1 | | < 0.00 | | < 0.00 |
| Scenario | 0.060 | 0.008 | 567 | | 6.85 | | < 0.00 | | < 0.00 |
| I(Scenario^2) | -0.017 | 0.002 | 567 | | -6.45 | | < 0.00 | | < 0.00 |
| Timepoint | 0.002 | 0.001 | 567 | | 1.55 | | 0.121 | | 0.604 |
| Scenario:Timepoint | -0.000 | 0.000 | 567 | | -1.14 | | 0.252 | | 1 |
